# Transcriptional program-based deciphering of the MET exon 14 skipping regulation network

**DOI:** 10.1101/2024.09.13.612820

**Authors:** Marie-José Truong, Geoffrey Pawlak, Jean-Pascal Meneboo, Shéhérazade Sebda, Marie Fernandes, Martin Figeac, Mohamed Elati, David Tulasne

## Abstract

The MET exon 14 skipping mutation (named METex14del) described in lung cancer leads to prolonged activation of signaling pathways and aberrant cell responses, but the link between HGF signaling and cell responses remains unclear. A putative regulatory network of influential regulators of target genes was constructed from the transcriptomes of lung cancer cell lines. Overlaying this reference network with transcriptomic data from METex14del-expressing cells, stimulated or not by HGF, revealed a major regulatory node consisting mainly of the transcription factors ETS1, FOSL1 and SMAD3. HGF activation of METex14del induced the phosphorylation of these master regulators and the expression of their predicted target genes in a RAS-ERK pathway-dependent manner. Furthermore, most of the transcription factors in the regulatory node are known regulators of epithelial-mesenchymal transition, consistent with their involvement in migration and invasion. New modeling with transcriptomic data from MEK inhibitor-treated METex14del cells validated the key role of RAS-ERK pathway regulators and their target genes in METex14del receptor activation. Thus, we report an original strategy to identify key transcriptional nodes associated with specific signaling pathways that may become novel therapeutic targets.

## Introduction

Receptor tyrosine kinases (RTKs) provide an interface between the extracellular and intracellular environments. Once activated by an extracellular growth factor, RTKs can activate intracellular signaling pathways that relay information within the cell, triggering various cell responses such as proliferation, differentiation, migration/invasion, and changes in the cell death/survival balance^1^. Signaling pathways transmit information to various cell compartments, including the nucleus, where regulation of gene expression is one of the most important mechanisms influencing cell responses ^2^. The transcriptional program, which can be deciphered by transcriptomic analyses, provides important information about biological responses, as gene expression profiles can be linked to key biological processes. However, how signaling pathways are integrated at the transcriptional level remains elusive.

In some cancers, RTK activation by genomic alterations can lead to oncogene addiction, in which cancer cell growth and/or survival is dependent on a single oncogene ^3^. In this situation, targeted therapies against RTKs, mainly tyrosine kinase inhibitors (TKIs), are effective and have profoundly improved patient care ^4^. In addicted cancer cells, overactive RTKs induce aberrant signaling leading to a deregulated transcriptional program. Thus, understanding oncogene addiction requires a deep understanding of signaling pathway integration at the transcriptional level. In addition, the efficacy of targeted therapies against RTKs is limited by primary or acquired resistance, often supported by activation of alternative signaling pathways that can bypass the targeted inhibition ^5^. In this situation, understanding signaling pathway integration at the transcriptional level is also an important issue.

MET, a member of the RTK family, is an emerging target in cancer. In lung cancer, the *MET* gene has been found to be mutated in approximately 3% of patients, with point mutations, deletions or insertions affecting the splice sites of exon 14. Such mutations result in in-frame skipping of this exon (METex14del) ^6,7^ and thus deletion of the juxtamembrane regulatory domain of the receptor ^8,9^.

Experiments using ectopic expression of METex14del or its reconstitution by genome editing in normal lung epithelial cells and cancer cell lines have shown that activated METex14del induces enhanced and sustained activation of downstream signaling pathways, including the RAS-ERK and PI3K-AKT pathways ^10–13^. In normal epithelial cells, METex14del activation depends on stimulation by HGF, its high-affinity ligand, and the mutant receptor can promote experimental tumor growth only in humanized mice expressing human HGF ^14^. Thus, in contrast to most RTK-activating mutations described to date, which result in ligand-independent RTK activation, MET exon 14 skipping does not eliminate MET receptor dependence on ligand stimulation. This suggests the existence of different activation states depending on ligand availability.

The juxtamembrane domain of MET contains several negative regulatory sites: (i) phosphorylated serine 985, involved in downregulation of MET tyrosine kinase activity ^15,16^, (ii) phosphorylated tyrosine 1003, involved in recruitment of the E3 ubiquitin ligase CBL and thus important for MET degradation ^17–19^, (iii) aspartic acid 1002, a caspase cleavage site involved in regulation of the death/survival balance ^20–23^. Downregulation of MET signaling appears to be mediated primarily by the CBL binding site ^10^.

Activated METex14del can also induce a much more complex transcriptional program than its WT counterpart. Gene ontology analyses are consistent with the migration and invasion responses induced by METex14del ^14^. However, this approach based on mRNA levels does not take into account the complexity of gene regulation, which involves the coordinated action of multiple transcription factors and co-factors (TF/co-TFs) and the influence of signaling pathways on their regulation.

The recently developed software solution CoRegNet ^24^, which can be visualized with the interactive tool Cytoscape Widget ^25^, is based on expression correlations between known cooperative TFs/co-TFs and target genes extracted from transcriptomic data and on described interactions between these regulators. CoRegNet has already proven its ability in complex organisms to identify 12 microglia-specific transcriptional regulators ^26^, 10 key regulators driving the transition in non-alcoholic liver disease^27^, and five master regulators specific to rheumatoid arthritis synovial fibroblasts ^28^.

Here we constructed a lung cancer-specific regulatory network based on the aggregation of extensive transcriptomic data from lung cancer cell lines. This network was then used to dynamically model the network of METex14del, stimulated or not by HGF. A “dialogue” between biological experiments and modeling confirmed the robustness of the proposed network and allowed an integrative view of METex14del signaling from its downstream signaling pathways to gene expression, which in turn regulates cell responses.

## Results

### Transcriptome-based modeling of the putative regulatory network of HGF-stimulated METex14del

Using the H-LICORN algorithm and the CoRegNet package ^24^, we constructed a lung cancer-specific regulatory network using transcriptomic data from 206 lung cancer cell lines from the CCLE database. The network was then enriched with supporting regulatory information such as TF binding sites, ChIP-seq data, and protein interactions found in the CoRegNet-embedded databases ^29–31^. The lung cancer-specific regulatory network consists of 502 key regulators that potentially regulate 4143 target genes with 18053 regulator/target gene regulatory interactions. The key regulators are connected by 2014 described protein-protein interactions (p <1e-130) and 61,272 transcription factor binding sites on their target genes (p <1e-300). A unique feature of CoRegNet is the implementation of transcriptional influence measures to estimate the activity state of a key regulator in a given sample. The influence of a regulator was calculated if at least five target genes were regulated with the highest correlation factor.

Normalized expression levels of differentially expressed genes (DEGs) from transcriptomic datasets of 16HBE MET WT and 16HBE METex14del in the presence and absence of HGF ^11^ were mapped onto this lung cancer regulatory network to analyze the effect of HGF stimulation on cells expressing MET WT (Supplementary Fig. S1 a and b) or METex14del (Supplementary Fig. S1 c, d). The transcriptional program of MET WT differs from that of METex14del, and two regulatory states are observed depending on whether cells are simulated by HGF or not. This suggests that the influence of key regulators differs between the two conditions. In these dynamic networks, the key regulators are referred to as “differentially influential regulators” (DIRs). The radius of the circle is proportional to the number of target genes of that DIR and the color of the circle represents its positive (red) or negative (blue) influence. A closer look at the model for HGF-stimulated 16HBE cells expressing METex14del reveals a large node of positive influence, which we have named “major regulatory node”. It is formed by six positively influencing interconnected DIRs (ETS1, FOSL1, SMAD3, HMGA2, CCND1 and RUNX1) with a large number of target genes (large circles) (Fig. 1a and Supplementary Fig. S1d). The representation by mean difference in influence (Fig. 1b) and the high value of the mean group influence (Supplementary Table. S2) confirm the activation state of these six major DIRs. Furthermore, the relationship between ETS1, SMAD3 and FOSL1 is characterized by protein-protein interactions between them (blue lines) and by the fact that the transcription factors ETS1 and SMAD3 have binding sites on each other’s genes and FOSL1 has a binding site for SMAD3 (red arrows). Notably, based on the mRNA levels for these six regulators in the transcriptomic data, only FOSL1 and HMGA2 appeared to be significantly upregulated under HGF stimulation (Fig. 1c), but by quantitative RT-PCR, all six DIRs showed significant up-regulation of their mRNA-level expression in the presence of HGF, with a stronger up-regulation for FOSL1 and HMGA2 (Fig. 1d). Taken together, this approach incorporating transcriptomics, important connections between regulators, and transcriptional influence calculations revealed a set of regulators putatively involved in the regulatory network of the HGF-stimulated METex14del receptor.

**Fig. 1.**
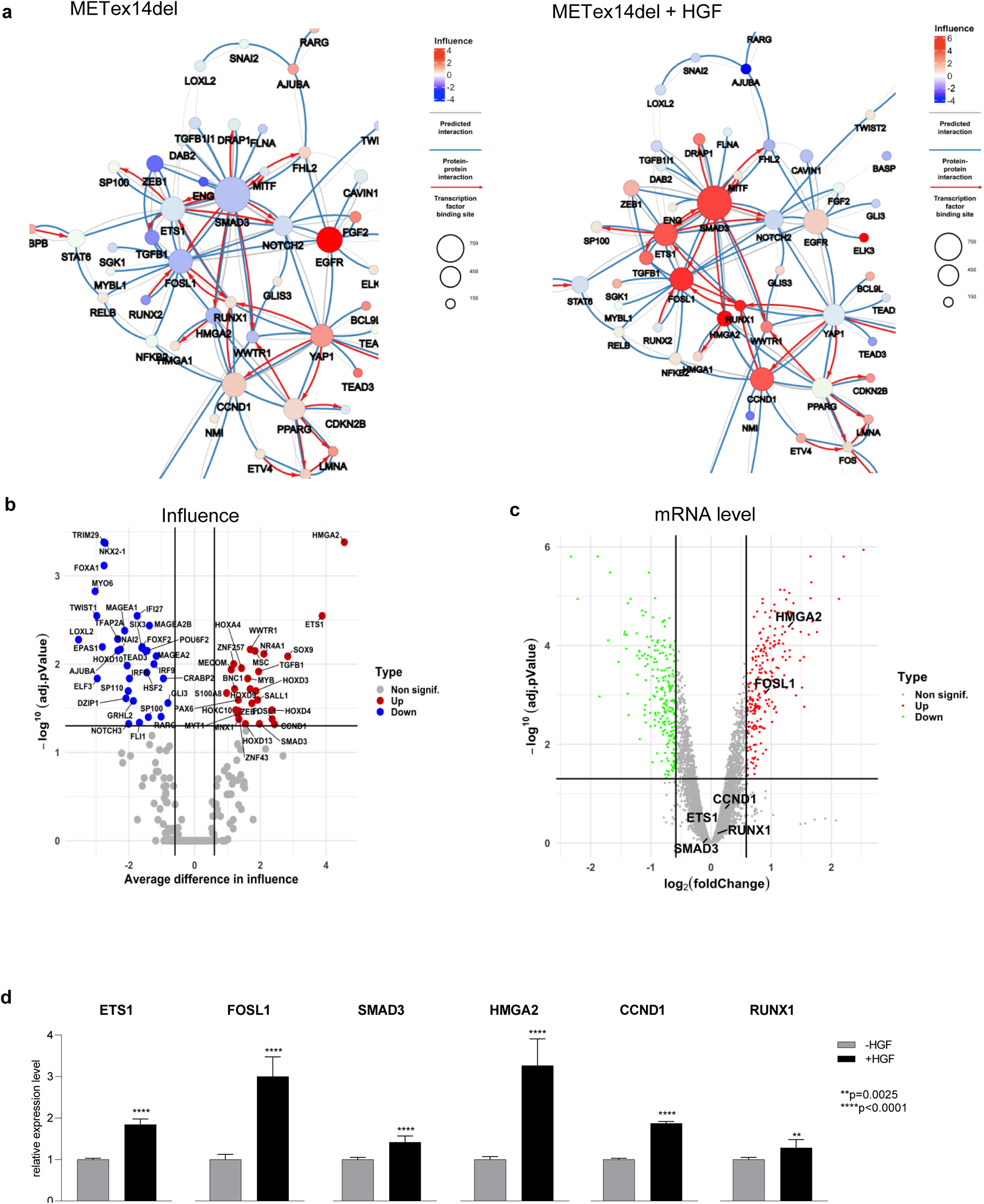
Regulatory network of METex14del in response to HGF. **a** Focus on the largest cluster of the co-regulatory network, showing the influence of TFs on METex14del cells in the presence and absence of HGF. Circles represent DIRs and the radius of the circle is proportional to the number of target genes regulated by the DIR. Co-regulatory interactions between DIRs are indicated: protein-protein interactions with published evidence (blue lines), transcriptional regulation interactions with published evidence (red arrows), and interactions defined only by the h-LICORN algorithm (gray lines). **b** DIRS were visualized in a volcano plot from interference network analysis: average difference in influence vs log10 transformed adj. P-value (regulators with an adj. p-value < 0.05 and an influence difference greater/smaller than 0.5 were considered up-/down-influenced (red/green). **c** DIRS were visualized in a volcano plot from transcriptomic analysis: log_2_ fold change vs p-value of differential mRNA-level expression (genes with an absolute adj. p-value < 0.05 and a fold change greater/smaller than 1.5 were considered upregulated/downregulated (red/green)). **d** mRNA-level expression of the six main influent DIRs present in the putative regulatory node, in the presence and absence of HGF (triplicates of *n*=3 independent experiments). Significance was determined by unpaired one-tailed *t*-test with Welch’s correction and data are expressed as mean *±* S.D.

### Evidence for a regulatory relationship between ETS1, FOSL1, and SMAD3 and their predicted target genes

We then focused on the three most important associated DIRs: ETS1, FOSL1, and SMAD3. Their influence was calculated according to their potential to regulate the expression of differentially expressed genes (DEGs). The transcriptional regulatory network of METex14del constructed with the microarray datasets allowed to visualize the putative influence of three DIRs, ETS1, FOSL1 and SMAD3 on their putative target genes divided into positive and negative genes (Fig. 2a). The 50 most regulated targets are shown in a heatmap (Fig. 2b), and the complete list of regulated targets by specific or combined DIRs was compiled (Supplementary Table. S3). A panel of ten targets up-regulated and two targets down-regulated under HGF stimulation, tested by RT-qPCR for expression at the mRNA-level, confirmed the predicted differential expression (Fig. 3a). Similar results were obtained for eight additional targets (six up-regulated and two down-regulated) (Supplementary Fig. S2). To assess the regulatory relationships between the three DIRs and their targets, ETS1, FOSL1, and SMAD3 were silenced by RNA interference, separately or in combination, and the expression of the original 12 targets was measured. The efficacy of each silencing was confirmed at the mRNA level by RT-qPCR (Fig. 3b) and at the protein level by Western blotting (Fig. 3c), under stimulation or not by HGF. Among the HGF upregulated target genes, eight (VIM, NOG, SERPINE2, SERPINA1, PTX3, CHGB, ABCA1 and LETM2) showed reduced expression in the presence of the three silencers, alone or in combination. The effects of ETS1 silencing on PTX3 expression and FOSL1 silencing on LETM2 expression were particularly pronounced (Fig. 3d). Reduced expression of SH2D5 was observed with either the ETS1 or FOSL1 silencer used alone and with the three silencers combined. Reduced expression of DCBLD2 was observed with either FOSL1 or SMAD3 used alone and with the three silencers combined. Of the two downregulated genes, IRS1 showed restored expression only in the presence of the siRNA combination and SLC15A3 also in the presence of the FOSL1 silencer alone (Fig. 3d). Taken together, these data show that all target genes tested are indeed regulated by at least one predicted DIR. Furthermore, with the exception of SLC15A3, which appears to be regulated only by FOSL1, all of the targets studied are regulated by two or three of these TFs. This indicates that most of the identified target genes are co-regulated by TFs of the node. Overall, these experimental data confirm the modeled co-regulatory network and reveal new regulatory relationships between master regulators and targets.

**Fig. 2.**
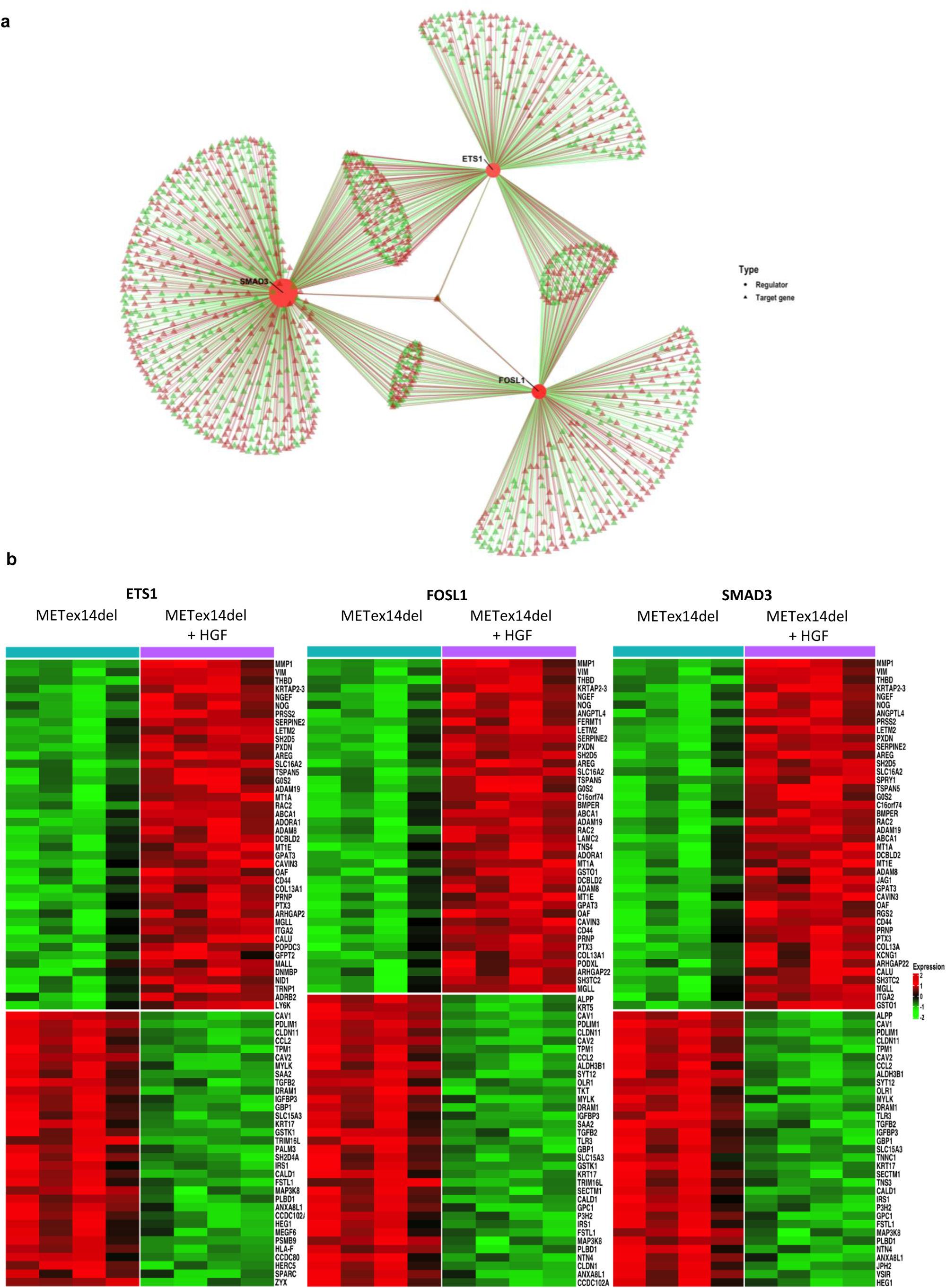
Identification of differentially expressed target genes potentially regulated by ETS1, FOSL1, and SMAD3. **a** Representations of the transcriptional regulatory network of ETS1, FOSL1 and SMAD3 with their putative influence on positively (red triangle) and negatively (green triangle) regulated DEGs. **b** Heatmaps showing putative ETS1, FOSL1 and SMAD3 target genes based on METex14del vs HGF-activated METex14del (adj p-value <0.05 and absolute fold change >1.5) (*n*=4 for each condition). Colors indicate high (red) and low (green) relative expression levels. **b** The mRNA-level expression of selected target genes up- or down-regulated in response to HGF was determined by RT-qPCR (triplicates of *n*=3 independent experiments). Significance was determined by unpaired one-tailed *t*-test with Welch’s correction, and data are expressed as mean *±* S.D.

**Fig. 3.**
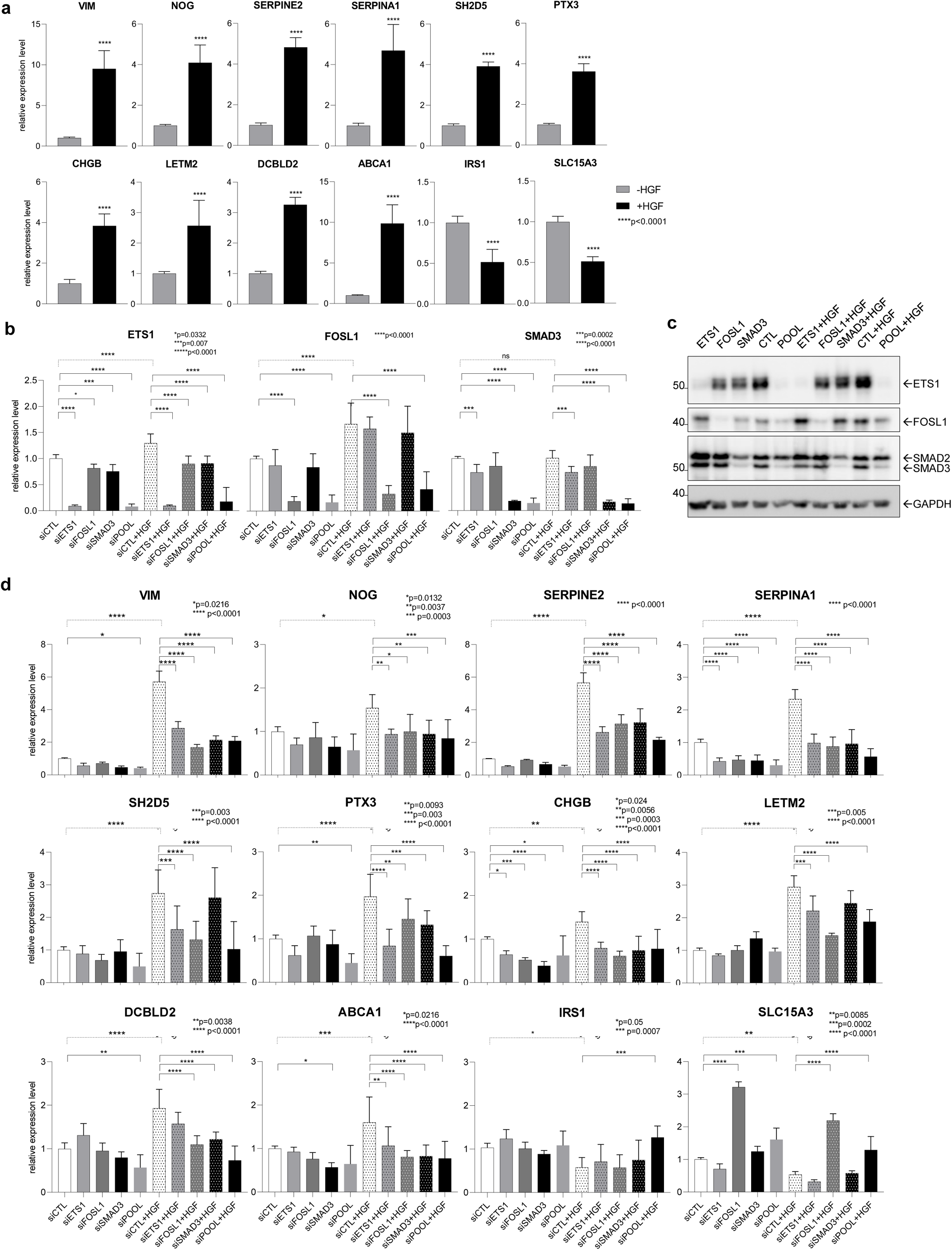
Effects of ETS1, FOSL1 and SMAD3 silencing on the expression of their predicted target genes. **a** The mRNA-level expression of selected target genes up- or down-regulated in response to HGF was determined by RT-qPCR (triplicates of *n*=3 independent experiments). Significance was determined by unpaired one-tailed *t*-test with Welch’s correction and data are expressed as mean *±* S.D. **b, c** The silencing efficacy of each specific siRNA (siETS1, siFOSL1, siSMAD3), used individually or in combination (siPOOL) was assessed by (**b**) RT-qPCR for mRNA-level expression (triplicates of *n*=3 independent experiments) and (**c**) by Western blotting for protein-level expression (representative experiment). **d** mRNA-level expression of up- and down-regulated target genes induced by knockdown of ETS1, FOSL1 and SMAD3 expression in 16HBE METex14del cells (triplicates of *n*=2, 3 or 4 independent experiments). Significance was determined by one-way ANOVA test and data are expressed as mean *±* S.D. in panels (**b**) and (**d**).

### Silencing of ETS1, FOSL1 and SMAD3 slightly affects wound healing and scattering

We then determined how HGF-induced biological responses are affected by silencing of the three TFs. HGF-stimulated METex14del cells show a migratory response consistent with the described transcriptional program^11^. Silencing of ETS1, FOSL1 or SMAD3 alone did not affect wound healing, but combined silencing slightly reduced both basal and HGF-stimulated responses (Fig. 4a). Similarly, in the presence of Matrigel, combined knockdown of all three TFs reduced HGF-induced wound invasion (Fig. 4b). HGF did not increase cell proliferation, but ETS1 knockdown enhanced this process both in the presence and absence of HGF (Fig. 4c). In addition, ETS1 knockdown appeared to increase HGF-induced scattering, whereas FOSL1 and SMAD3 knockdown decreased it (Fig. 4d). These results show that combined knockdown of the three TFs slightly reduces HGF-induced migration, invasion and scattering.

**Fig. 4.**
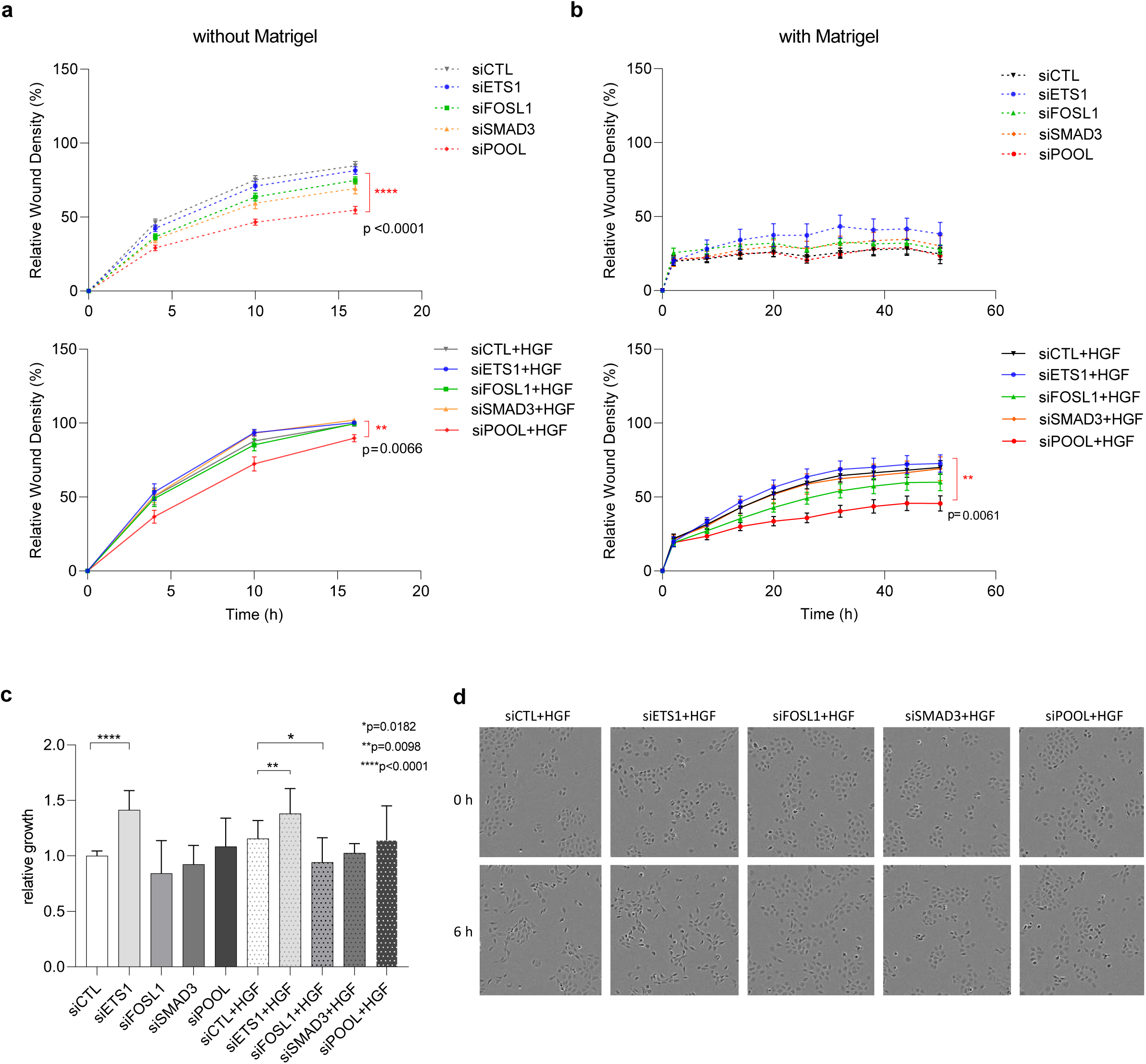
Effect of ETS1, FOSL1, and SMAD3 silencing on HGF-induced migration, invasion, proliferation and scattering. Forty-eight hours after transfection with siRNAs targeting ETS1, FOSL1 and SMAD3 alone or in combination, wound healing was monitored without (**a**) or with (**b**) Matrigel in the presence and absence of HGF (triplicate of *n*=4 independent experiments). In all graphs, only statistically significant differences between the negative control (siCTL) and the applied treatment (an siRNA used alone or pooled with the other siRNAs, siPOOL) in the presence or absence of HGF are indicated. Wound healing data are expressed as mean *±* S.E.M. and significance was determined by two-way ANOVA test. **c** The effect of siRNA on proliferation was determined by Alamar blue staining after 16 hours of stimulation with HGF and relative growth was expressed as relative mean fluorescence (6 replicates of *n*=4 independent experiments). Proliferation data are expressed as mean *±* S.D. and significance was determined by one-way ANOVA test. **d** Images of the effect of silencing on cell scattering were captured at different time points using the Incucyte system and representative images from baseline and 6 hours after stimulation with HGF are shown.

### The RAS-ERK signaling pathway controls the phosphorylation of ETS1, FOSL1, and SMAD3, as well as cellular wound healing

Since silencing of transcription factors has little effect on known biological functions induced by HGF, we investigated the signaling pathways that might act upstream of these TFs to understand how they might act together in the regulatory node. First, for all master regulators in MET WT and METex14del cells, we heat mapped the top 50 differentially expressed target genes upon HGF stimulation (Fig. 5a). Gene ontology (GO) enrichment analysis of these genes revealed significant overrepresentation of annotations such as epithelial-mesenchymal transition (EMT) (12.2%, p-adj=3.8E-20), KRAS signaling up (8.8%, p-adj=1.7E-08), positive regulation of locomotion (6.4%, p-adj=5.4E-12), cell junction organization (8.4%, p-adj=4.1E-19), apical junction (8.2%, p-adj=6.5E-07), or signaling by receptor tyrosine kinase (8.2%, p-adj=1.1E-08), all of which are consistent with cell responses induced by HGF in 16HBE cells (Fig. 5b and Supplementary Table. S4). Given our previous study showing that HGF/MET signaling induces activation of ETS1 through ERK-dependent phosphorylation of its threonine 38 (T38) ^35,36^, we examined the induction of ETS1 phosphorylation and also analyzed the phosphorylation of the other two master regulators, FOSL1 and SMAD3. As expected, HGF stimulation was found to induce ETS1 phosphorylation (at T38) in 16HBE METex14del cells, an effect detected up to 8 h post-stimulation. No phosphorylation of this TF was detected in their WT counterparts (Fig. 5c). Similar sustained ETS1 phosphorylation by HGF was observed in ZORG cells (Fig. 5c) and H596 cells (Supplementary Fig. S3a) derived from NSCLC patients harboring the METex14del variant. A slight increase in ETS1 expression was observed in METex14del cells 8 hours after HGF stimulation (Fig. 5c). FOSL1 is known to be phosphorylated by ERK at serines 252 and 265 (S252 and S265) ^37^. Strong induction of FOSL1 expression and phosphorylation (S265) was found in our three cell models: 16HBE METex14del cells, ZORG cells, and H596 cells, peaking at 8 h post-simulation and persisting until 24 h (Fig. 5d and Supplementary Fig. S3b). The main regulator of SMAD3 is known to be the TGFβ receptor, which induces phosphorylation at serine residues 423 and 425 (S423, S425) ^38^. However, it has been shown that in response to TGFβ, ERK can also phosphorylate SMAD3 at three serines: S204, S208, and S213, and that this also contributes to SMAD3 activation ^39^. Here, we found that in response to HGF, SMAD3 was phosphorylated at S208 in 16HBE MET WT, 16HBE METex14del, ZORG, and H596 cells, with maximum phosphorylation at 30 minutes after stimulation (Fig. 5e and Supplementary Fig. S3c). Taken together, these results indicate that HGF stimulation of METex14del induces sustained signaling of the RAS-ERK pathway, leading to phosphorylation of ETS1, FOSL1, and SMAD3. Thus, this signaling pathway is a positive activator of the major regulatory node of the METex14del network.

**Fig. 5.**
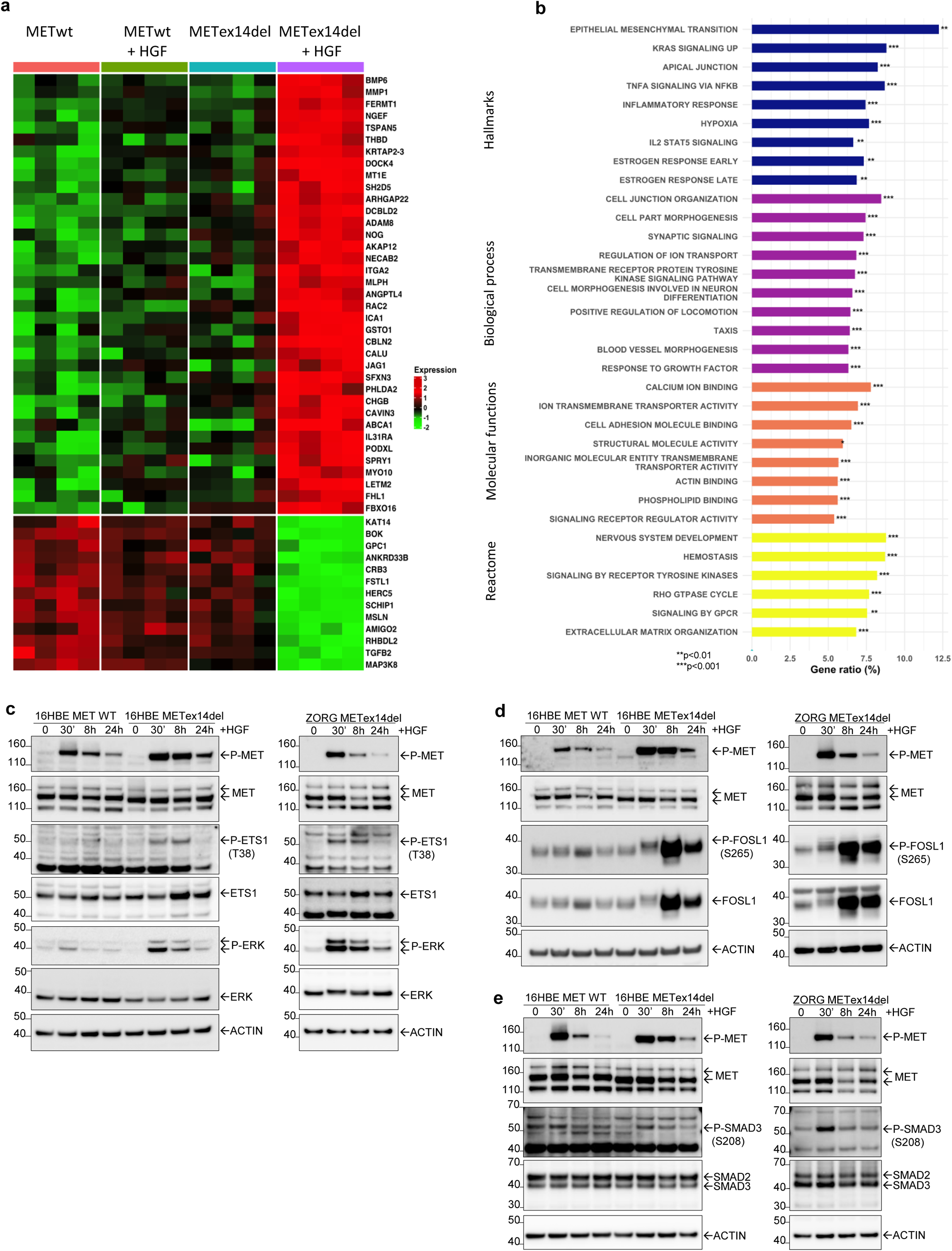
Gene ontology (GO) annotation of target genes induced by HGF activation of METex14del. **a** Heatmap of the 50 most significantly differentially regulated target genes in response to HGF (adj. p-value < 0.05 and absolute fold change >1.5) between indicated conditions (*n*=4 per condition). Colors indicate high (red) and low (green) levels of relative expression. **b** Shown are gene ratios (%) of GO enrichment for the differentially expressed target genes according to Hallmarks, Biological Process, Molecular Functions and Reactome annotations. Overrepresentation analysis was performed using the hypergeometric test: ** p-value < 0.01, ***p-value < 0.001. **c, d, e** The 16HBE cells expressing MET WT or METex14del and ZORG cells derived from lung cancer patients expressing METex14del were stimulated with HGF in a time course experiment. Expression and activation of (**c**) ETS1 and its form phosphorylated at T38 (P-ETS1), (**d**) FOSL1 and its form phosphorylated at S265 (P-FOSL1), and (**e**) SMAD3 and its form phosphorylated at S208 (P-SMAD3) were analyzed by Western blotting (representative results).

To confirm this signaling mechanism, HGF-stimulated cells were treated with trametinib, a well-known MEK inhibitor used in melanoma patients harboring the BRAF V600 mutation ^40^. In 16HBE METex14del and ZORG cells, HGF-induced phosphorylation of ETS1 at T38, FOSL1 at S265 and SMAD3 at S208 was reversed by the addition of either trametinib (Fig. 6a, b and c) or U0126, another MEK inhibitor (Supplementary Fig. S4a, b and c). These data confirm the involvement of the RAS-ERK signaling pathway in the activation of the key transcription factors ETS1, FOSL1 and SMAD3. The effect of trametinib on the biological responses of 16HBE METex14del cells was also investigated. Trametinib was found to inhibit HGF-induced wound healing both in the absence (Fig. 6d) and presence of Matrigel (Fig. 6e), although less effectively than the MET TKI capmatinib. Similar results were observed in cell scattering assays (Fig. 6f). Co-treatment with both capmatinib and trametinib resulted in more potent inhibition (Fig. 6d, e and f). Cell proliferation was only slightly affected by these inhibitors (Fig. 6g). Thus, consistent with the mild inhibition of METex14del-dependent motility and invasion observed upon knockdown of ETS1, FOSL1 and SMAD3, pharmacological inhibition of the RAS-ERK signaling pathway, which regulates the activity of these three transcription factors, dramatically inhibited these cell responses.

**Fig. 6.**
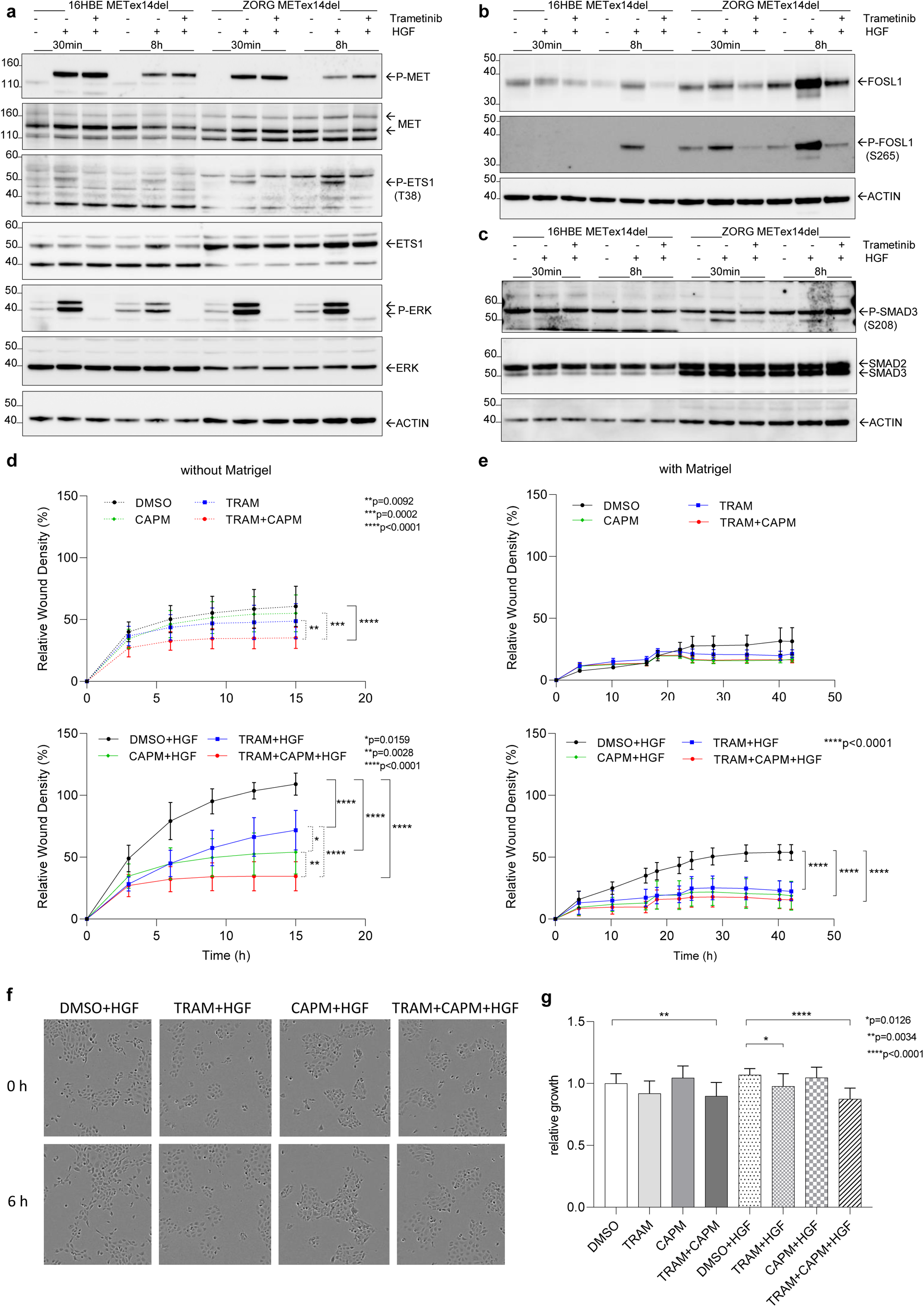
Effect of MEK inhibitor on ETS1, FOSL1 and SMAD3 phosphorylation and cell responses induced by HGF. The effect of trametinib (TRAM, a MEK inhibitor) on the expression of (**a**) P-ETS1/ETS1, (**b**) P-FOSL1/FOSL1 and (**c**) P-SMAD3/SMAD3 in 16HBE METex14del cells and ZORG cells stimulated or not with HGF was determined by Western blotting (representative results). The effects of trametinib and capmatinib (CAPM, a MET inhibitor), alone and in combination, on cell wound healing were evaluated without (**d**) and with (**e**) Matrigel (six replicates of *n*=3 independent experiments); (**f**) scattering in 16HBE METex14del cells stimulated or not with HGF (representative photo of baseline and 6 hours after inhibition under HGF) and (**g**) proliferation (six replicates of *n*=4 independent experiments). In all graphs, only statistically significant differences between the negative control without HGF (DMSO) and the different treatments (individual or combined inhibitors) in the presence or absence of HGF are indicated. Wound healing data in panels (**d,e**) are expressed as mean *±* S.E.M. and significance was determined by two-way ANOVA test. In panel (**g**), proliferation data are expressed as mean *±* S.D. and significance was determined by one-way ANOVA test.

### Involvement of the RAS-ERK signaling pathway in the positive regulation of the METex14del regulatory network

A new 3’RNA-seq transcriptomic analysis was performed on 16HBE METex14del cells stimulated or not with HGF and treated or not with capmatinib or trametinib. Our previous transcriptomic results had shown that HGF stimulation induced many DEGs. In these new experiments, capmatinib treatment restored the transcriptional program to the unstimulated state and trametinib treatment induced a drastic change in DEGs under both basal and HGF-stimulated conditions (Fig. 7a). To verify the effect of these inhibitors on the activation of DIRs of the regulatory network, we mapped the normalized gene expression levels of DEGs from our RNAseq datasets onto the lung cancer-specific reference network (Fig. 7b and Supplementary Figure S5 a, b, c, d, e, and f). HGF treatment alone resulted in an activated network profile similar to Fig. 1a, with activation of the same six positively influential TFs of the major regulatory node: ETS1, FOSL1, SMAD3, HMGA2, CCND1 and RUNX1 (Fig. 7b). As expected, capmatinib completely inhibited their positive influence (Supplementary Table S5). Interestingly, a stronger reversal of the influence of these DIRs was observed after trametinib treatment, suggesting that the RAS-ERK signaling pathway is the main activator of the entire regulatory node in the lung cancer network. What’s more, when expression of the previously analyzed regulated target genes of ETS1, FOSL1, and SMAD3 were plotted in a box-and-whisker plot (Fig. 7c), most of them (VIM, NOG, SERPINE2, SERPINA1, SH2D5, CHGB, LETM2, DCBLD2, IRS1, and SLC15A3) showed the expected up- or down-regulation under HGF stimulation, which was reversed by inhibitor treatments. The exception was ABCA1, which showed only weak activation (Fig. 7c). As expected, direct blockade of the MET receptor with capmatinib did not allow HGF/MET to induce stimulation, and blocking only the RAS-ERK signaling pathway with trametinib prevented the activation of some specific HGF-activated DIRs and the corresponding target genes. Overall, these new data confirm that activation of the main node of the co-regulatory network modeled in lung cancer, involving master influencers and their targets, is mainly mediated by the RAS-ERK signaling pathway.

**Fig. 7.**
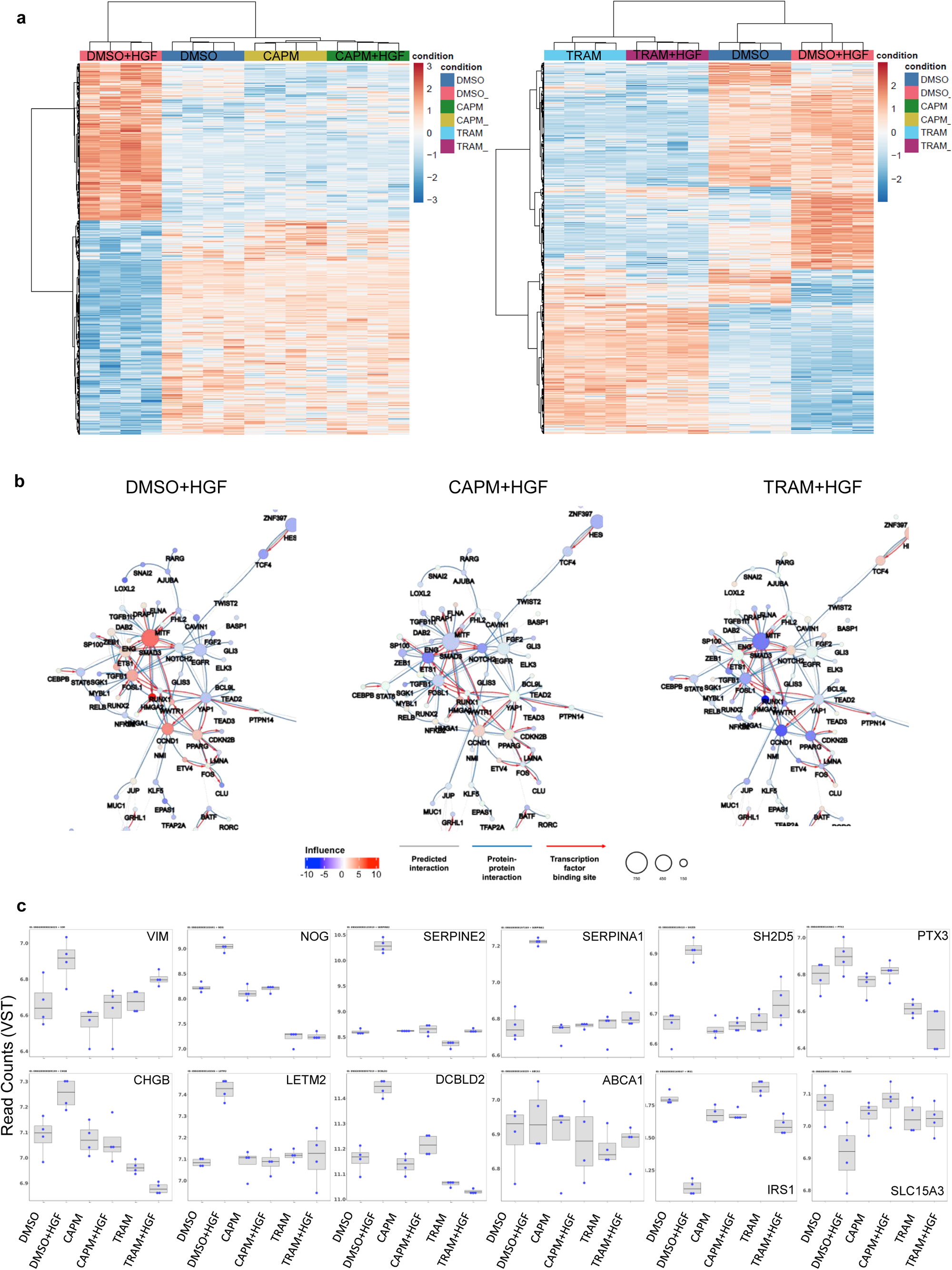
Effect of MET and MEK inhibitors on the transcriptional program induced by HGF and on the predicted regulatory network. **a** Heatmaps of differentially expressed genes (adj. p-value < 0.05 and absolute fold change > 1.5) in 16HBE METex14del cells treated with capmatinib (CAPM) or trametinib (TRAM) and stimulated or not with HGF are shown (*n*=4 per condition). **b** Zooms on the major regulatory node of the co-regulatory network of METex14del cells in the presence and absence of inhibitors (CAPM, TRAM) upon stimulation with HGF. The radius of the circle is proportional to the number of target genes of the respective DIR. Co-regulatory interactions between transcription factors are indicated: protein-protein interactions with published evidence (blue line), transcriptional regulation interactions with published evidence (red arrow), and interactions defined only by the h-LICORN algorithm (gray line). **c** Expression of selected target genes regulated by ETS1, FOSL1, SMAD3 are shown in box-and-whisker plots. The box plot shows the 25th, 50^th^, and 75th percentiles, while the blue dots show the variant stabilization transformation (VST) of gene expression for each sample (*n*=4 per condition). The median is indicated by the line across the box.

## Discussion

It has been shown that disease phenotypes, including those related to disease progression and response to therapy, are maintained by small groups of TFs and co-TFs ^41,42^. This justifies research efforts to identify these factors. Here, we have used a reverse engineering approach to infer the first lung cancer-specific gene regulatory network from the lung cancer cell dataset without any a priori knowledge. We have used this network to identify key regulators (essentially transcription factors) that putatively influence the expression of target genes involved in HGF-induced responses in cells expressing an activated form of MET. Importantly, we find that our METex14del regulatory network can distinguish two distinct states of cell activation: HGF-stimulated or not.

There are several different bioinformatic methods to infer gene regulatory networks from high-throughput data, thus enabling the discovery of disease-driving genes and/or pathways. Successful approaches to date include those (e.g., ARACNe, LICORN, GENIE3) that allow reverse-engineered construction of context-specific networks and which can be further enriched by integrating interaction evidence (protein-protein interactions and/or transcriptional regulation) ^43^. The bioinformatics approach proposed in this study is original, mainly in terms of (i) incorporating cooperativity between co-regulators into the model ^44,45^, which makes it closer to biological reality than most other approaches, (ii) considering the influence of regulators by evaluating the expression of target genes rather than the expression of the regulators themselves, with the aim of detecting master regulators ^45^, (iii) the fact that reference-based analysis shifts data interpretation from unsupervised to supervised, allowing accumulated information from previous experiments (here 206 untreated lung cell lines) to help interpret new data.

The case of the regulatory network of HGF-stimulated METex14del presented here demonstrates the potential of the above strategy to improve the analysis and interpretation of small transcriptomic datasets. Indeed, with the regulatory network of HGF-stimulated METex14del, we have uncovered the putative positive influence of an important regulatory node, consisting in particular of three interconnected transcription factors: ETS1, FOLSL1 and SMAD3. The connections between DIRs were established by their known functional relationships, including protein-protein interactions (red arrows), and also by their predicted commonly regulated target genes (gray lines). Several studies have already documented interactions between these DIRs: co-regulation of gene expression by ETS1 and SMAD3 ^46–48^ and the existence of active transcriptional complexes containing them ^49^; an association between FOSL1 and SMAD3 leading to co-regulation of target genes has also been demonstrated ^50^, as has a potential cooperation between ETS1, SMAD3 and AP1 to regulate the angiopoietin-like 4 gene^51^.

Since the network predicts a strong positive influence of these transcription factors, we assume that they are activated by HGF stimulation. Transcription factors can be activated by several mechanisms, including expression, phosphorylation state, localization, and interaction with co-activators. In our model, we found that FOSL1 expression is strongly increased under HGF stimulation, but ETS1 and SMAD3 expression is only weakly increased. This suggests a different activation mechanism for the latter two, independent of protein expression. Using antibodies recognizing ETS1 phosphorylated at Thr38, FOSL1 phosphorylated at Ser265, and SMAD3 phosphorylated at Ser208, we confirmed that HGF stimulation of the METex14del receptor activates all three TFs by inducing their phosphorylation. Note that we and others have previously shown that these phosphorylations of ETS1 and FOSL1, known to be ERK-dependent ^52–55^, can be induced by HGF stimulation in cells expressing the WT form of MET ^56,57^. Interestingly, phosphorylation of FOSL1 is known to prevent its ubiquitination and thus lead to its stabilization ^55^, a fact consistent with the strong increase in FOSL1 expression upon HGF stimulation. SMAD3 is part of the canonical TGFβ signaling pathway ^58^. The TGFβ receptor activated by its ligand can phosphorylate SMAD2 and SMAD3 on C-terminal serine residues ^59^, allowing them to associate with SMAD4 and regulate gene expression after nuclear translocation. SMAD3 can also be phosphorylated by ERK and other kinases on serine residues within the linker domain ^60^. These SMAD2 and SMAD3 phosphorylations were initially found to inhibit TGFβ-dependent signaling by inhibiting nuclear translocation ^61^, but other studies have shown that RAS-ERK-dependent phosphorylation of SMAD2 and SMAD3, in response or not to TGFβ, can promote SMAD-dependent transcription and cell transformation, particularly in the context of cancer cells. This suggests that SMAD linker phosphorylation plays different roles depending on the cell context ^62^. Consistent with our model, we show here that the activated RAS-ERK signaling pathway induced by HGF-stimulated METex14del can also phosphorylate SMAD3, presumably regulating its transcriptional activity. Consistent with our findings, it has been shown that HGF and TGFβ can cooperate to induce JNK-dependent phosphorylation of SMAD2 and SMAD3 in the linker region, in normal cells of gastric origin ^60^. In addition, it has recently been shown in A549 lung cancer cells expressing METex14del (as a result of genome editing) that SMAD2 is phosphorylated and able to associate with MET. Cell invasion and migration and experimental metastatic spread have been shown to be inhibited by SMAD2 silencing ^13^. Thus, both our data and data from the literature identify TGFβ-independent activation of SMAD signaling as a relay of METex14del activity.

To further investigate the transcriptional program, we confirmed that HGF can regulate many of the ETS1, FOSL and SMAD3 target genes as predicted and that silencing of the three TFs can restore the expression of most of the affected target genes. This suggests that the regulation of these genes is supported by multiple transcription factors of the regulatory node identified here. Taken together, these results validate the robustness of the model and highlight an important regulatory node involving three transcription factors acting together to regulate target genes.

Functionally, we also show that inhibition/silencing of ETS1, FOSL1 or SMAD3 alone does not alter METex14del-induced cell motility and invasion, but that significant inhibition can be achieved when all three TFs are inhibited together. This suggests that these three regulators work together to regulate migration and invasion and that additional transcription factors may be involved. It is worth noting that all three TFs belong to a family of transcription factors, the closest members of which are ETS2, FOS/JUN and SMAD2. The fact that these TFs were not identified as master regulators in our METex14del model suggests that these close family members may only be activated when their homologs are silenced, creating a compensatory mechanism of action.

Epithelial-to-mesenchymal transition (EMT) is an important step in cell migration and invasion, particularly through reduction of intercellular adhesion, loss of apical-basal polarity, and gain of motility ^63^. Several growth factors can regulate EMT, the most important being TGFβ, but also HGF and EGF. Interestingly, most of the transcription factors in the identified regulatory node are known to activate EMT. In particular, SMAD3 is the key regulator of TGFβ signaling capable of inducing EMT ^64^. Several studies have identified FOSL1 as a regulator of EMT through rapid induction of morphological changes and both down- and up-regulation of EMT markers ^65^. ETS1 has been shown to be a regulator of EMT, particularly in the context of TGFβ stimulation, with cross-regulation with the EMT transcription factors ZEB, SNAIL, and TWIST ^66^. In our study, however, the role of ETS1 may be broader, as its knockdown slightly increases cell proliferation and cell scattering. Although the functional involvement of HMGA2 was not investigated in this study, it has been shown to induce EMT, particularly in lung cancer cells ^67,68^. It is worth noting that ZEB1 and TWIST2, key transcription factors in EMT, are also associated with the regulatory node. This suggests their involvement, which should be investigated in functional studies. Consistently, several confirmed target genes of ETS1, FOLSL1, and SMAD3 are involved in EMT, including vimentin and integrin subunit alpha 2 (ITGA2) ^69,70^. Overall, most of the transcription factors in this regulatory node are known regulators of EMT, consistent with their involvement in migration and invasion.

Since ETS1, FOSL1 and SMAD3 are regulated by a common signaling pathway, the RAS-ERK signaling pathway, we sought to determine how inhibition of this pathway affects the regulatory network. First, we show that a clinically used TKI directed against MET can restore the basal transcriptomic program regulated by HGF-stimulated METex14del, as visualized by both the DEGs heatmap and the regulatory network. Furthermore, a MEK inhibitor, which is mainly used clinically for the treatment of melanoma can profoundly alter gene expression, under both basal and HGF-stimulated conditions. Interpretation of these transcriptomic data through the regulatory network reveals that within the main regulatory node, most regulatory influences are inhibited by this treatment. This suggests that the RAS-ERK signaling pathway positively regulates the influence of the entire node of the network. We further confirmed the central role of RAS-ERK signaling with functional data showing that METex14del-induced migration and invasion were abolished by the MEK inhibitor.

Taken together, our transcriptomics-based regulatory network and the functional experiments performed to support our interpretations highlight a number of key transcription factors that act together to regulate the biological responses induced by METex14del. In particular, inhibition of a single TF of the master node did not perturb the biological responses, and only simultaneous knockdown of three of them resulted in significant inhibition. Furthermore, inhibition or restoration of target gene expression required knockdown of two or three of them. Thus, the regulation of genes by multiple TFs forming an interconnected node could ensure the robustness of the network, with the knockdown of one regulator being insufficient to perturb the biological outcome. To go further, we interpreted “influence” as post-transcriptional activation, i.e. phosphorylation, of the transcription factors, rather than simply increased expression. This led us to identify the RAS-ERK pathway as an important signaling pathway that can positively regulate both the overall major regulatory node and the associated biological responses. Importantly, unlike the TFs, this pathway is readily targeted by pharmacological approaches, opening the way to novel therapeutic strategies, particularly in the context of incomplete response to MET TKIs in patients harboring METex14del mutations.

## Methods

### Cell Lines

The non-tumorigenic 16HBE14o-(RRID:CVCL_0112) (16HBE) cell line, developed by immortalization of primary human bronchial epithelial cells with SV40 large T antigen, was a kind gift from Pr. Dieter Gruenert ^71^. The parental 16HBE cell line (MET WT) and derived cells (METex14del) ^11^ were maintained in GIBCO MEM (Thermo Scientific) supplemented with 10% heat-inactivated FBS (Sigma-Aldrich). The ZORG cell line was derived from a pleural effusion of a MET-TKI resistant patient^72^. The H596 and Hs746T cell lines (HTB-135TM and HTB178TM) were obtained from the American Type Culture Collection (Manassas, Virginia). Hs746T cells were maintained in DMEM (Thermo Scientific) and H596 and ZORG cells were maintained in RPMI1640. All media were supplemented with 10% FBS. All cells were routinely tested for mycoplasma using MycoAlertTM (Lonza). Cells were cultured at 37°C under a humidified controlled atmosphere of 5% CO2 in air. For MET activation experiments, cells were serum starved in serum-free medium for 16 h after overnight adherence and before HGF stimulation or inhibitor treatment (pretreatment with 1 µM inhibitor for 3 h before activation by HGF for 30 min, 8 h or 24 h).

### Reagents and RNA interference

HGF (Miltenyi Biotec B.V. & Co. KG) was dissolved in 5% BSA in PBS. U0126 (Promega), trametinib (MedChemTronica), and capmatinib (Selleck Chemicals) were prepared in dimethyl sulfoxide (DMSO). The ETS1 (VHS40612 and VHS40614), FOSL1 (HSS188462 and s15585), and SMAD3 (VHS41114 and VHS41111) siRNAs and the Stealth siRNA negative control Lo GC (12935200) and Silencer® Select Negative Control #1 siRNA (#4390843) were purchased from Thermo Fischer Scientific. Transfections were performed using Lipofectamine™ RNAiMAX (Thermo Fisher Scientific) and siRNA at 50 nM per well according to the manufacturer’s protocolCells were used for further experiments 48 hours after transfection.

### Immunoblotting

The Nucleospin RNA/Protein Kit (Macherey-Nagel) was used to extract RNA and also proteins from siRNA validation experiments (Fig. 3b). All other proteins were extracted with RIPA buffer supplemented with protease and phosphatase inhibitor cocktail (1% aprotinin, 1 mM PMSF, 1 µM leupeptine, 1 mM Na3VO4, 20 mM β-glycerophosphate), sonicated for 20 s, and centrifuged at 1500g for 15 min. Protein in the supernatant was quantified using the BCA protein assay kit (Thermo Scientific). Equal amounts of protein from cell lysates were resolved on a NuPAGE Bis-Tris polyacrylamide gel (Invitrogen) and transferred to Immobilon-P polyvinylidene fluoride (PVDF) membranes (Millipore). After blocking with casein buffer (0.2% casein in PBS containing 0.1% Tween20), the membranes were sequentially incubated with the indicated primary antibodies and HRP-conjugated secondary antibodies. Chemiluminescence signals were detected using SuperSignal West Dura or Femto (Thermo Scientific). Image analysis was performed using ImageQuant LAS3000 (GE Healthcare, US). Primary antibodies were provided by Cell Signaling Technology, Inc. (Danvers, MA) for anti-MET (C-terminal, #3148), anti-phospho-MET (Tyr1234/Tyr1235, #3126), anti-phospho-ERK1/2 (Thr202/Tyr204; #9106), anti-ETS1 (D808A, #14069), anti-phospho-FRA1 (Ser265, #3880), and anti-SMAD2/3 (D7G7, #8685) (all at 1:1000 dilution); Santa Cruz Biotechnology, Inc. (Dallas, Texas, USA) for anti-ERK2 (sc-154), anti-FRA1 (C12, sc-28310), anti-GAPDH (sc-32233, 1:10,000 dilution); Abcam Limited (Cambridge, UK) for anti-phospho-ETS1 (T38, #ab59179) and anti-phospho-SMAD3 (S208, #ab138659). The following peroxidase-conjugated secondary antibodies were used at 1:10,000 dilution: anti-mouse (#115-035-146) and anti-rabbit (#711-035-152), purchased from Jackson ImmunoResearch (Cambridge Shire, UK).

### RT-qPCR analysis

After total RNA extraction using the Nucleospin RNA/Protein Kit (Macherey-Nagel) and quantification using NanoDrop™ 2000 (Thermo Scientific), cDNA was reverse transcribed using the High Capacity cDNA Reverse Transcriptase Kit (Invitrogen). Quantitative polymerase chain reaction (qPCR) was performed using Fast SYBRTM Green Master Mix (Applied Biosystems) on a QuantStudio3 (Applied BiosystemsTM, Thermo Fisher Scientific). All samples were performed in triplicate. Relative expression levels were calculated using the 2-ΔΔCt method. The level of each transcript was normalized to the β2m (beta-2 microglobulin) expression level. Primers used are listed in Supplementary Table. S1.

### Scratch wound healing assay

After 24 hours of attachment, cells (transfected or not) were starved with 0.1% FBS medium for 4 hours before scratch wounds were made using a 96-well WoundMaker (Essen BioScience) according to the manufacturer’s instructions. For the invasion assay, Matrigel matrix (growth factor reduced, Corning) was added to each well at 50 μL/well and allowed to solidify for 30 min in a 37°C CO2 incubator according to the manufacturer’s instructions. Cells were then treated with or without inhibitor (1 µM) in the presence or absence of HGF (20 ng/mL) (n=3 to 6 for each condition). Images of the wounds were captured automatically in the CO2 incubator using IncuCyte Zoom or SX5 software (Essen BioScience). Wound images were updated every 2 hours for the duration of the experiment. Data were analyzed for wound confluence and calculated using the IncuCyte software package (Essen BioScience).

### Proliferation and scattering cell imaging with Incucyte

After 24 h of attachment, cells (transfected or not) were starved for 4 h in 0.1% FBS medium before treatment or not with inhibitor (1 µM) in the presence or absence of HGF (20 ng/ml) for 16 h. They were then transferred to an IncuCyte incubator (n=6 for each condition). Cell growth and viability were quantified using AlamarBlue™ Cell Viability Reagent (Invitrogen) and fluorescence was measured using a Multiskan RC Microplate Reader (ThermoLabsystems) with 560/590 (ex/em) wavelength filter settings. Each result was expressed as the average optical density of four independent experiments. For scattering, images were acquired automatically in the CO2 incubator using the IncuCyte system (Essen BioScience). Wound images were updated every 2 hours for the duration of the experiment.

### Transcriptomic data source

The Cancer Cell Line Encyclopedia (CCLE) project ^73^ provides transcriptomic data for lung cancer cell lines downloaded from the Cancer Dependency Map portal (DepMap Broad (2024); https://depmap.org/portal.) using the depmap R package. This resulted in 206 cell lines, including 135 NSCLC cell lines, 50 SCLC cell lines and 21 others. Of the 206 cell lines, 105 were derived from primary tumors and 101 from metastatic tumors. Transcriptomic data from 16HBE cells expressing MET WT or METex14del, stimulated or not by HGF, are available in the Sequence Read Archive under accession number PRJNA764905 (GSE184514, raw data).

### Public data collection of regulatory elements and interactions

The Human Transcription Factor Catalog ^74^ and the TcoF-DB v2 database ^75^ provided a list of 2,375 regulators corresponding to 1,639 transcription factors (TFs) with experimentally validated DNA binding specificity and 752 transcription cofactors (TcoFs) with experimental validation information. Human transcription factor binding sites (TFBSs) were modeled using MotifDB R/Bioconductor (Shannon P, Richards M (2024). *MotifDb: An Annotated Collection of Protein-DNA Binding Sequence Motifs*. R package version 1.46.0. DOI: 10.18129/B9.bioc.MotifDb) and promoter sequences were searched using with the PWMEnrich R/Bioconductor package (Stojnic R, Diez D (2024). *PWMEnrich: PWM enrichment analysis*. R package version 4.40.0. DOI: 10.18129/B9.bioc.PWMEnrich). ChIP-seq data from the ChEA2 database ^76^ and the tftargets R package (https://github.com/slowkow/tftargets) and various databases (e.g. TRED, ITFP, ENCODE, BEDOPS, TRRUST) provided all regulatory evidence from TF to target gene, yielding a list of 2,256,674 TF-target interactions. The list of evidence of TF-TF cooperation, containing 1,257,053 protein-protein interactions (PPIs) from various PPI databases (e.g., BioGrid, HIPPIE, STRING, and HPRD) was obtained using the iRefR R package ^77^.

### Gene regulatory network inference

Gene regulatory networks were inferred using the CoRegNet Bioconductor package ^24^. Starting with transcriptomic data and a list of human regulators, CoRegNet uses the hLICORN algorithm ^78^ to capture regulatory interactions between regulators and target genes in four steps: (1) Transcriptomic data are discretized into values of -1, 0, and 1 according to the distribution per gene. Genes present in the transcriptomic dataset were partitioned into regulators and target genes, and only those with minimum support for non-zero values after discretization were retained. (2) An priori frequent itemset algorithm identified potential sets of co-activators and co-inhibitors from the list of regulators. (3) For each target gene, a list of candidate sets of co-activators and co-inhibitors (GRN) is selected according to a regulatory program (association rule) metric. (4) Each candidate GRN for each gene is scored based on the regression between the expression of regulators in the GRN set and the expression of the gene in question. For each gene, we retained the top ten candidate GRNs according to the R2 score. CoRegNet can then be used to select the best GRN for each gene based on the interaction evidence data. Evidence of regulation and cooperation was integrated into each candidate GRN using an integrative selection algorithm, resulting in an R2 score for each of the integrated datasets. The GRN with the highest integrated score was selected. CoRegNet constructs a coregulatory network from the obtained GRNs by establishing a cooperativity relationship between pairs of regulators, TFi and TFj, if they share at least five target genes and the relationship is significant (p =< 0.01) according to the Jaccard similarity coefficient formula: 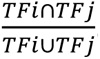.

### Quantification of regulatory influence signals

The regulatory network structure provides a set of genes that are activated or repressed under reference conditions. Based on this structure, we can capture the influence of each regulator, a latent signal of regulator activity in each sample based on its observed effect on downstream entities. For each regulator, Welch’s t-test was performed to compare the distribution of activated *(A^r^)* and repressed *(I^r^)* genes. The influence of a regulator r is calculated as follows

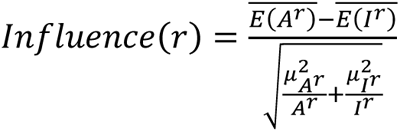

where *E(A^r^)* and *E(I^r^)* are expressions of the activated and repressed genes in the samples, respectively, 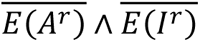 are their respective means, and μ*A^r^* and μ*I^r^* are their standard deviations. A regulator is active only if it activates A^r^ and represses I^r^ according to the expectations of the regulatory network model, resulting in a positive *t*-value in the Welch *t*-test. Higher values of *In*(*r*) indicate a higher level of activity of the regulator *r* in the sample. In this work, we calculated the *influence* of a regulator only if it activated or repressed at least five genes. Due to the strict thresholds used in network construction, some local gene networks could be filtered out, resulting in some regulators having activated or repressed target gene sets with fewer than five members. In such cases, we filled the smaller set (less than 5 target genes) with the target genes whose regulator had the highest R2 regression value. This allowed us to capture the *influence* values for a maximum number of regulators. In addition, for some regulators, expression values might be missing for some of the target genes in some of the tumor samples whose *influence* we wanted to assess. In this case, the missing expression values were estimated using the LatNet method ^79^, with expression levels in cell lines as a reference dataset. In some (rare) cases, it was not possible to estimate the expression of some target genes in a particular sample, which could prevent the calculation of the *Influence* of a particular regulator. In such cases, the median of the *influence* values obtained for the regulator in the other samples was used to replace the missing value.

### Identification of group-specific regulators and target genes

Gene expression analysis was performed and differentially influential regulators (DIRs) were identified using linear models for microarray data (LIMMA) R package ^80^, and *p*-values were adjusted using the Benjamini-Hochberg method. Gene expression analysis was performed on the normalized gene expression data set, and differential influence analysis was performed on the influence data. Regulators with an adj. p-value < 0.05 were considered DIRs.

### Functional enrichment analysis of differentially expressed genes (DEGs)

Gene Ontology (GO) enrichment analysis was performed using overrepresentation analysis methods with the R packages clusterProfiler ^81^ and enrichR ^82^. Ontologies with an adj. p-value<0.05 and gene counts of more than five genes were examined. The Hallmarks, Biological Processes Database (MSigDB), Molecular Function and REACTOME pathway databases were searched for overrepresented terms using the R package msigdbr ^83^.

### Sequencing library construction and library sequencing

ERCC spike-in was added to each sample as a control. 3’RNA-Seq library generation (QuantSeq 3’ mRNA-Seq Library Prep Kit FWD) was initiated by oligo dT priming with 200ng of total RNA. The primers already contain Illumina compatible linker sequences. After first strand synthesis the RNA is degraded and second strand synthesis is initiated by random priming and a DNA polymerase. The random primer also contains 5’ Illumina-compatible linker sequences and Unique Molecular Identifiers (UMIs). After obtaining the double-stranded cDNA library, the library is purified with magnetic beads and amplified (14 cycles). During the library amplification, the barcodes (UDI) and sequences required for cluster generation (index i7 in 3’ and index i5 in 5’) are introduced via Illumina-compatible linker sequences. The final libraries are purified and loaded onto a High Sensitivity DNA Chip controlled by an *Agilent Bioanalyzer 2100*. Library concentration and size distribution are checked.Each library is pooled equimolarly and the final pool is sequenced on a *Nova 6000 instrument* (Illumina) with 100 cycles of chemistry.

### Data analysis for transcriptome data

To remove poor quality regions and poly(A) from the reads, we used the *fastp* program. We used a quality score threshold of 20 and removed reads shorter than 25 pb. Read alignments were performed using the *STAR* program with the human genome reference (GRCh38) and reference gene annotations (*Ensembl*). The UMI (Unique Molecular Index) allowed us to reduce errors and bias in the PCR were processed using *fastp* and *umi tools*. Based on the read alignments and the UMI, we counted the number of molecules per gene using *FeatureCount (from 3.73M to 11.9M molecules, 6.82M on average)*. Other programs used for read quality control and workflow were Q*ualimap*, *fastp*, *FastQC* and *MultiQC*. Differential gene expression of RNA-seq was performed using the *R/Bioconductor* package *DESeq2*. The cut-off for differentially expressed genes was adj. p-value< 0.05 and absolute fold change >1.5. RNAseq data were deposited in the Sequence Read Archive under accession number PRJNA1123466.

### Statistics

All results except Network inference (DEGs, DIRs, …) and transcriptomic analysis are expressed as mean ± S.E.M. or S.D. for the indicated number of independent experiments. Data were analyzed using GraphPad Prism® 9 (San Diego, CA, USA).

## Acknowledgements

This work was supported by the Cancéropôle Nord-Ouest (Emergent 2022, https://canceropole-nordouest.org/), INCa (NETMET-2022-74012, https://www.e-cancer.fr/Institut-national-du-cancer/), and La Ligue Contre le Cancer (MET exoCancer-2021-74012, https://www.ligue-cancer.net/).

## Author contributions

M-J.T. and DT directed the project and contributed to funding acquisition; conceived and designed the experiments and interpreted the main results; wrote the manuscript with the help of all authors.

M.Fe, JP.M., S.S., D.T., M.Fi. contributed to the transcriptomic analysis (microarrays) M-J.T., JP.M., S.S., D.T., M.Fi. contributed to the transcriptomic analysis (RNAseq) G.P., and M.E. conceived and designed the co-regulatory networks.

All authors provided critical feedback and helped shape the research, discussed the results, and commented on the manuscript.

## Data available

Micro-array transcriptomic data from 16HBE cells expressing MET WT and METex14del stimulated or not by HGF (GSE184514, raw data are available in the Sequence Read Archive under accession number PRJNA764905). The RNAseq data from 16HBE with or without treatment (CAPM/TRAM) were deposit in the Sequence Read Archive (SRA) under accession number PRJNA1123466.

