## Supplemental Figures for "Transcriptional program-based deciphering of the MET exon 14 skipping regulation network"

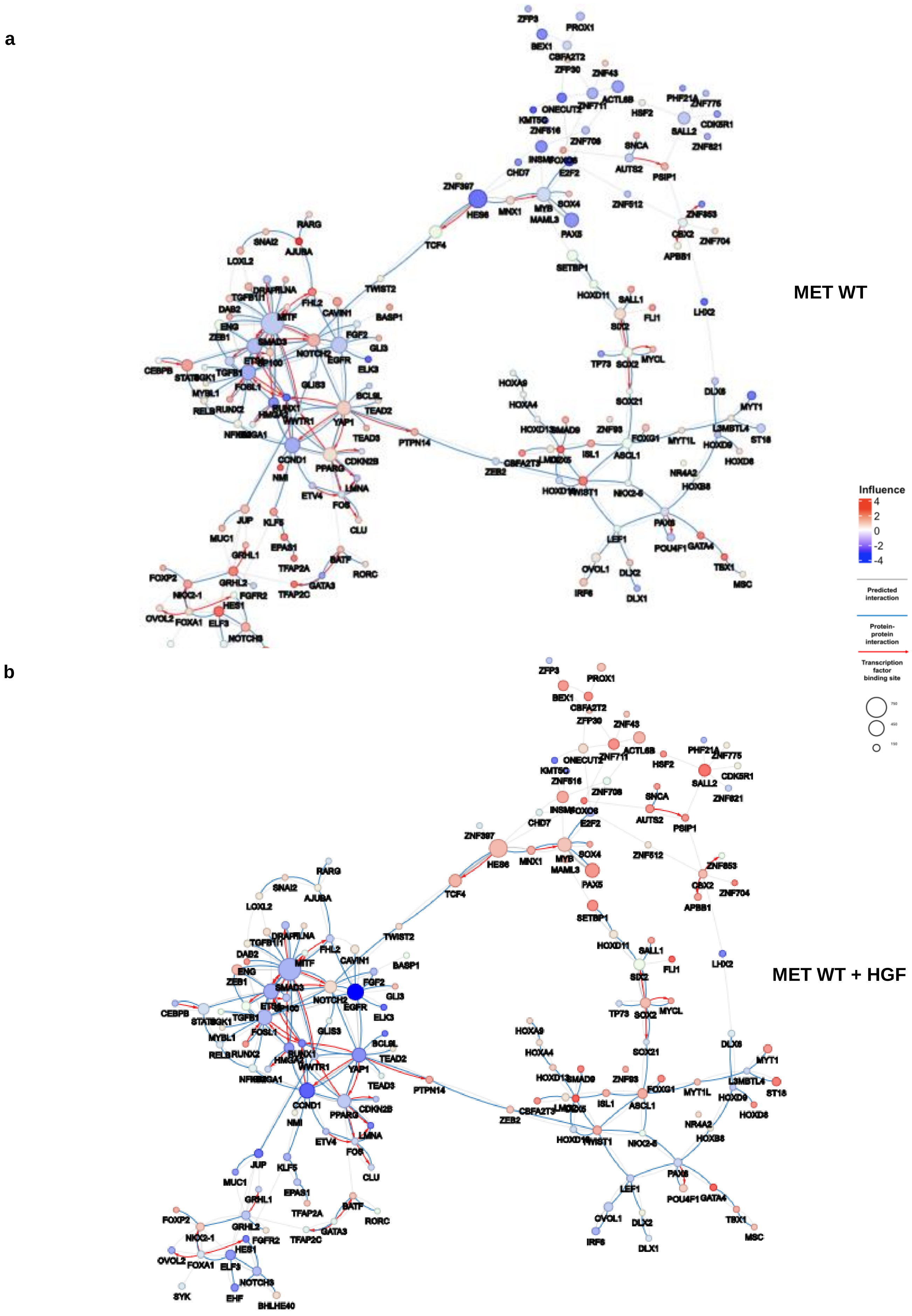
a
MET WT
b
MET WT + HGF
Influence
4
2
0
-2
-4
Predicted interaction
Protein-protein interaction
Transcription factor binding site
750
450
150

c

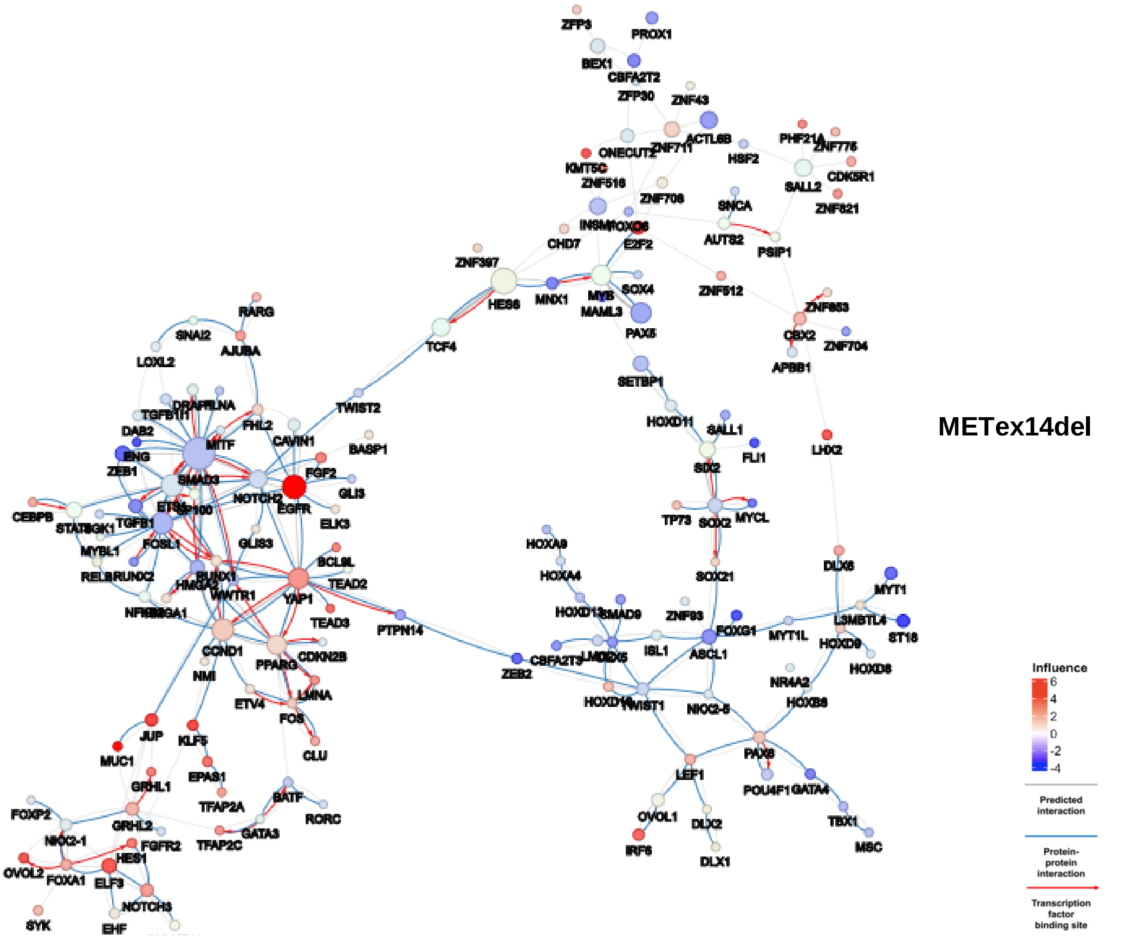
METex14del
Influence
6
4
2
0
-2
-4
Predicted interaction
Protein-protein interaction
Transcription factor binding site
750
450
150
ZFP3
PROX1
BEX1
CBFA2T2
ZFP30
ZNF43
ACTL6B
ONECUT2
ZNF711
KMT5C
ZNF516
ZNF706
HSF2
PHF21A
ZNF775
CDK5R1
SALL2
ZNF821
SNCA
AUTS2
PSIP1
INSM1
FOXO6
E2F2
CHD7
ZNF397
HES6
MNX1
MYB
SOX4
MAML3
PAX5
ZNF512
ZNF853
CBX2
ZNF704
APBB1
TCF4
SETBP1
HOXD11
SALL1
FLI1
SIX2
LHX2
TWIST2
RARG
SNAI2
AJUBA
LOXL2
DRAP1
FLNA
TGFB1I1
FHL2
DAB2
MITF
CAVIN1
BASP1
ENG
ZEB1
SMAD3
FGF2
GLI3
NOTCH2
EGFR
ELK3
CEBPB
STAT5
SGK1
TGFB1
SP100
ETS1
FOSL1
GLIS3
MYBL1
BCL9L
RELB
RUNX2
RUNX1
HMGA2
TEAD2
NFIB
HMGA1
WWTR1
YAP1
TEAD3
PTPN14
CCND1
CDKN2B
PPARG
NMI
LMNA
ETV4
FOS
CLU
JUP
KLF5
MUC1
EPAS1
GRHL1
TFAP2A
BATF
RORC
FOXP2
GRHL2
GATA3
NKX2-1
FGFR2
TFAP2C
HES1
OVOL2
FOXA1
ELF3
NOTCH3
SYK
EHF
TP73
SOX2
MYCL
SOX21
HOXA9
HOXA4
HOXD13
SMAD9
ZNF83
FOXG1
MYT1L
ISL1
ASCL1
CBFA2T3
LMO2
ZEB2
HOXD10
TWIST1
NKX2-5
DLX6
MYT1
L3MBTL4
ST18
HOXD9
HOXD8
NR4A2
HOXB8
PAX8
LEF1
POU4F1
GATA4
OVOL1
DLX2
TBX1
IRF6
DLX1
MSC

d

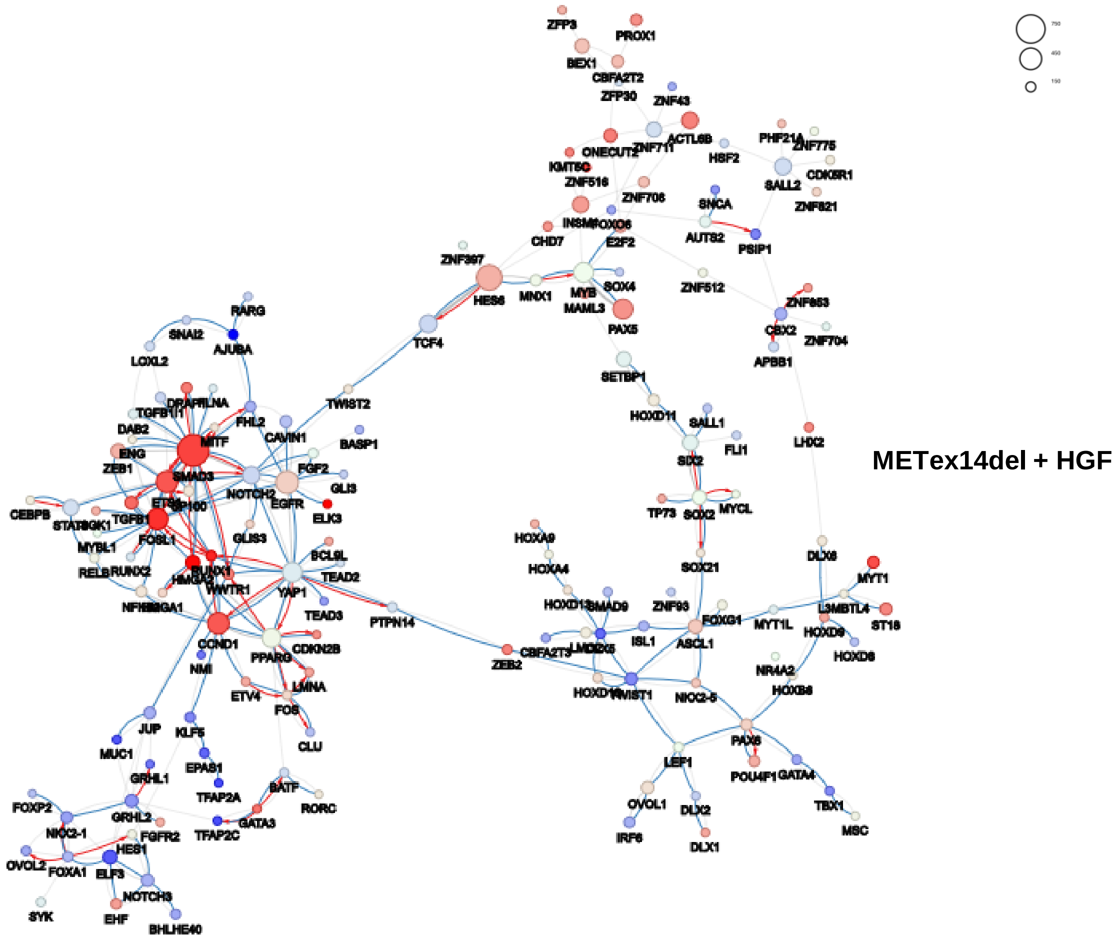
METex14del + HGF
ZFP3
PROX1
BEX1
CBFA2T2
ZFP30
ZNF43
ACTL6B
ONECUT2
ZNF711
KMT5C
ZNF516
ZNF706
HSF2
PHF21A
ZNF775
CDK5R1
SALL2
ZNF821
SNCA
AUTS2
PSIP1
INSM1
FOXO6
E2F2
CHD7
ZNF397
HES6
MNX1
MYB
SOX4
MAML3
PAX5
ZNF512
ZNF853
CBX2
ZNF704
APBB1
TCF4
SETBP1
HOXD11
SALL1
FLI1
SIX2
LHX2
TWIST2
RARG
SNAI2
AJUBA
LOXL2
DRAP1
FLNA
TGFB1I1
FHL2
DAB2
MITF
CAVIN1
BASP1
ENG
ZEB1
SMAD3
FGF2
GLI3
NOTCH2
EGFR
ELK3
CEBPB
STAT5
SGK1
TGFB1
SP100
ETS1
FOSL1
GLIS3
MYBL1
BCL9L
RELB
RUNX2
RUNX1
HMGA2
TEAD2
NFIB
HMGA1
WWTR1
YAP1
TEAD3
PTPN14
CCND1
CDKN2B
PPARG
NMI
LMNA
ETV4
FOS
CLU
JUP
KLF5
MUC1
EPAS1
GRHL1
TFAP2A
BATF
RORC
FOXP2
GRHL2
GATA3
NKX2-1
FGFR2
TFAP2C
HES1
OVOL2
FOXA1
ELF3
NOTCH3
SYK
EHF
BHLHE40
TP73
SOX2
MYCL
SOX21
HOXA9
HOXA4
HOXD13
SMAD9
ZNF83
FOXG1
MYT1L
ISL1
ASCL1
CBFA2T3
LMO2
ZEB2
HOXD10
TWIST1
NKX2-5
DLX6
MYT1
L3MBTL4
ST18
HOXD9
HOXD8
NR4A2
HOXB8
PAX8
LEF1
POU4F1
GATA4
OVOL1
DLX2
TBX1
IRF6
DLX1
MSC

**Supplementary Fig. S1 Regulation network of MET WT and METex14del in response to HGF.** Co-regulatory network showing the influence of TFs on MET WT in the absence (**a**) or presence of HGF (**b**) and on METex14del in the absence (**c**) or presence of HGF (**d**). Circles represent DIRs and the radius of the circle is proportional to the number of target genes regulated by the DIR. Co-regulatory interactions between DIRs are indicated: protein-protein interactions with published evidence (blue lines), transcriptional regulation interactions with published evidence (red arrows), and interactions defined by the h-LICORN algorithm only (gray lines).

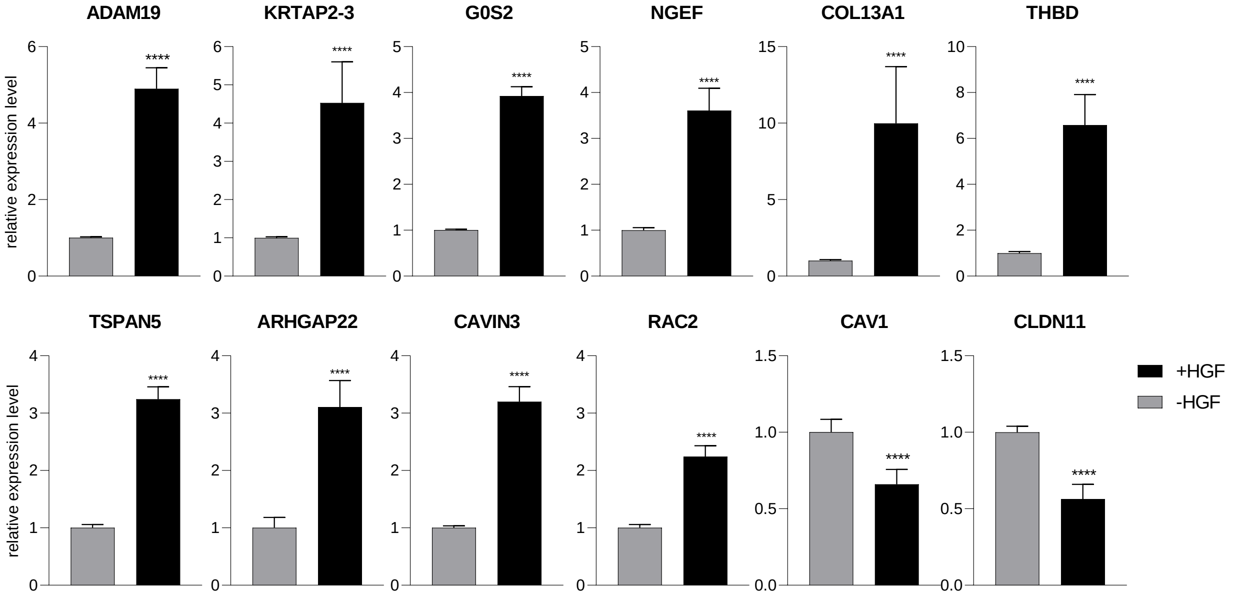


**Supplementary Fig. S2 Effects of HGF on the expression of target genes potentially regulated by ETS1, FOSL1 and SMAD3.** The mRNA-level expression of additional target genes up- or down-regulated in response to HGF was determined by RT-qPCR (triplicates of *n*=4 independent experiments). Significance was determined by unpaired one-tailed *t*-test with Welch's correction and data are expressed as mean ± S.D.

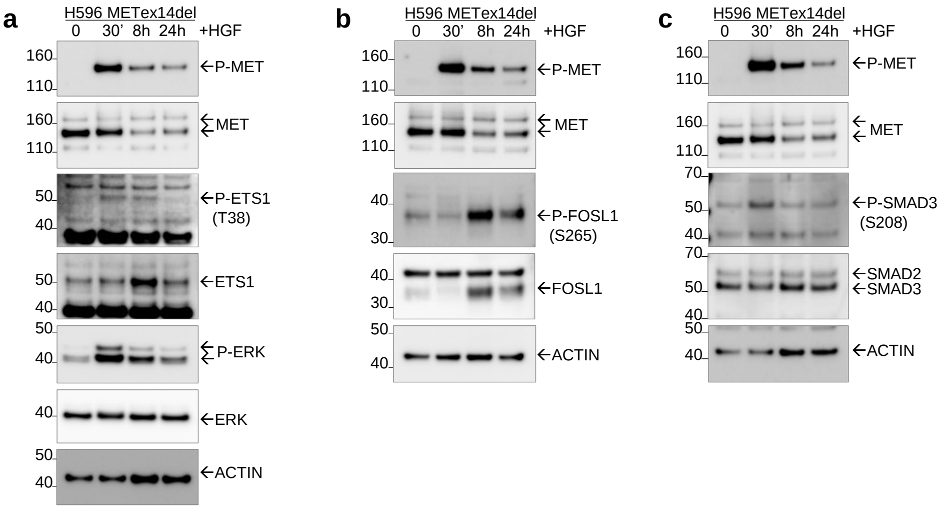


**Supplementary Fig. S3 Time course effect of HGF on expression and phosphorylation of ETS1, FOSL1 and SMAD3 in H596 cells.** Expression and activation of (**a**) ETS1 and its phosphorylated form at T38 (P-ETS1), (**b**) FOSL1 and its phosphorylated form at S265 (P-FOSL1), and (**c**) SMAD3 and its phosphorylated form at S208 (P-SMAD3) were analyzed by Western blot (representative results).

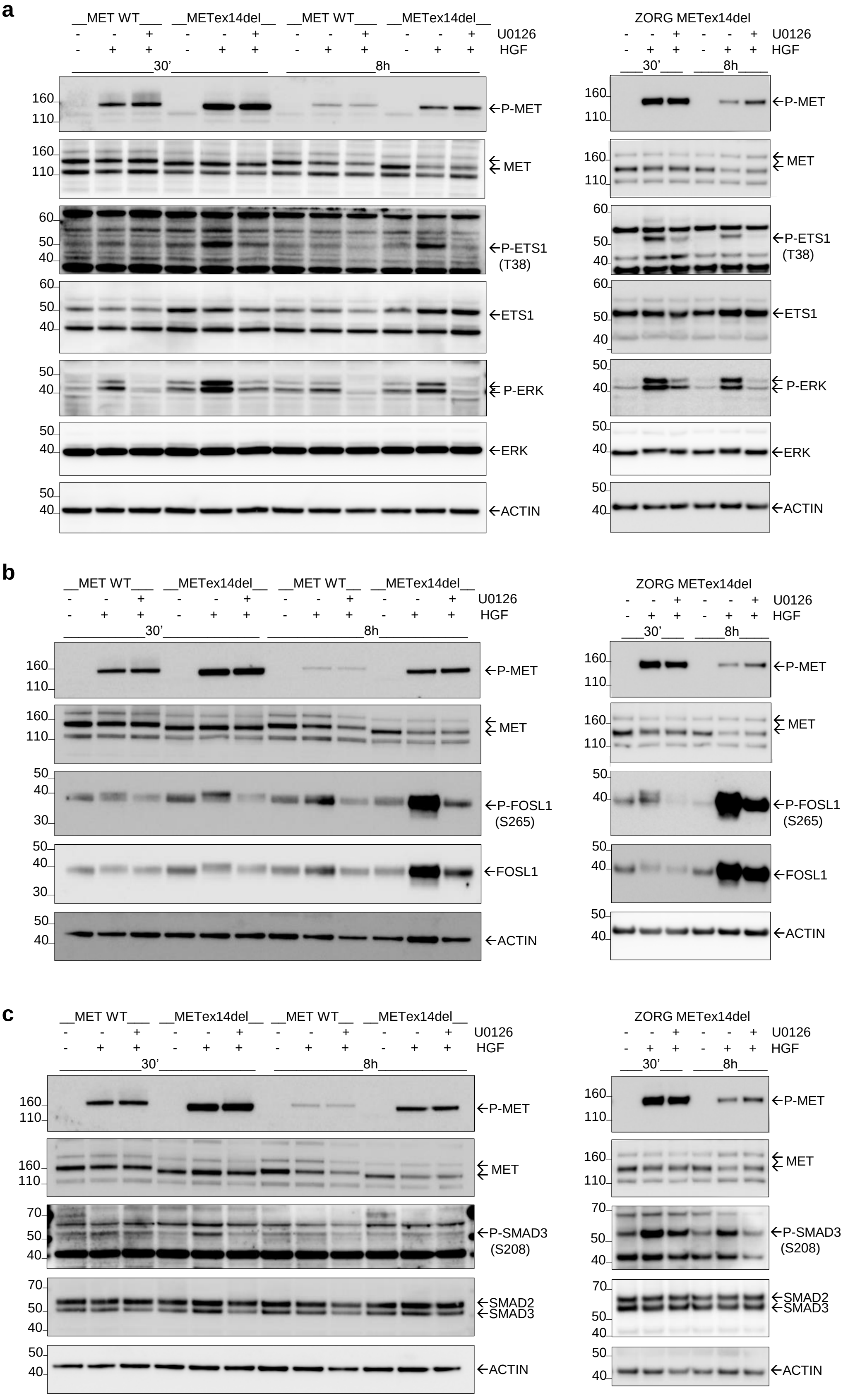


**Supplementary Fig. S4 Effect of U0126 on ETS1, FOSL1 and SMAD3 expression and phosphorylation.** U0126 (10µM) was added 3 hours before stimulation with HGF. The expression of (**a**) P-ETS1/ ETS1, (**b**) P-FOSL1/FOSL1 and (**c**) P-SMAD3/SMAD3 was analyzed in 16HBE MET WT and METex14del cells and ZORG cells stimulated or not with HGF (representative results) as indicated.

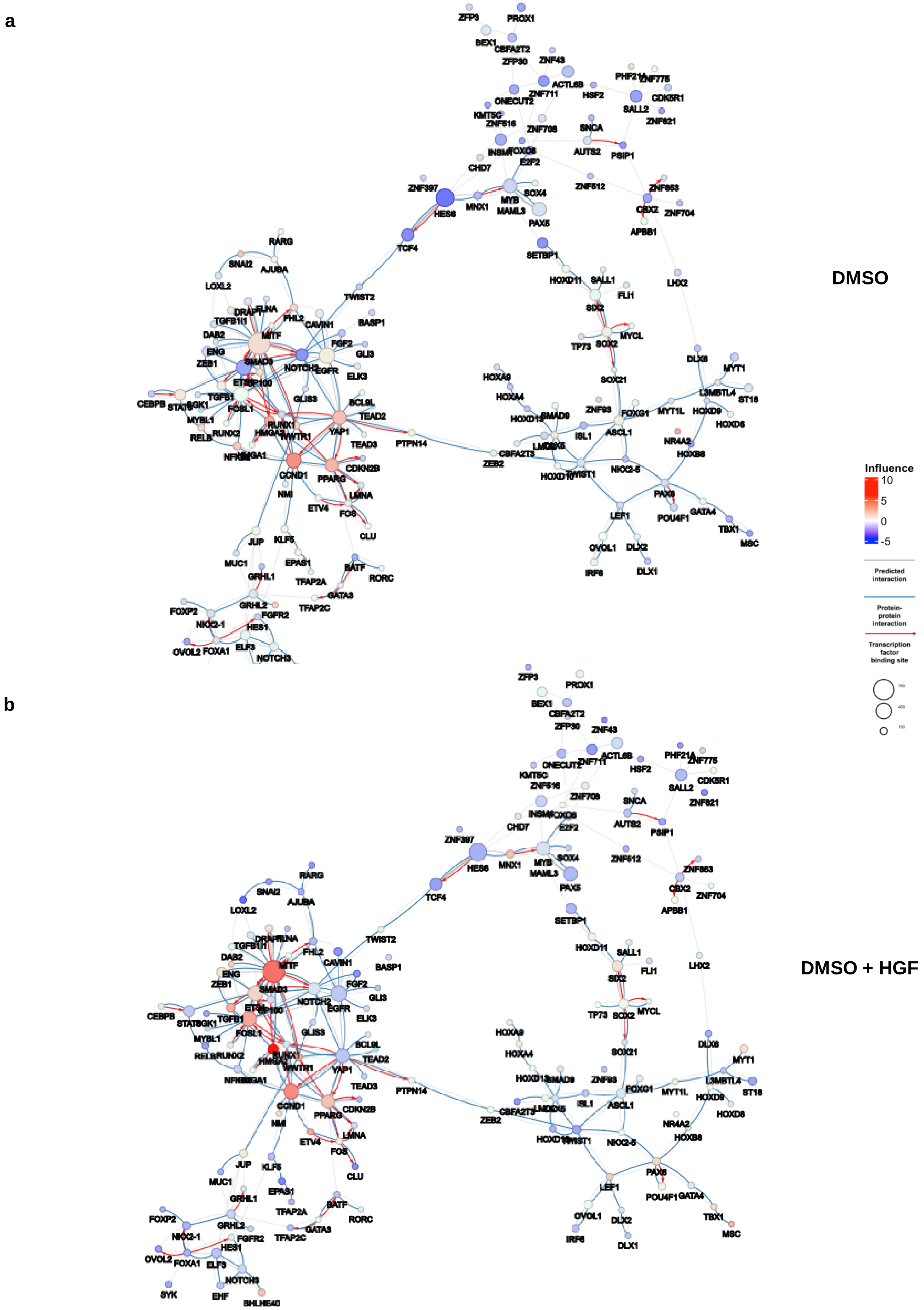
a
DMSO
b
DMSO + HGF
Influence
10
5
0
-5
Predicted interaction
Protein-protein interaction
Transcription factor binding site
ZFP3
PROX1
BEX1
CBFA2T2
ZFP30
ZNF43
ACTL6B
ZNF711
HSF2
PHF21A
ZNF775
CDK5R1
SALL2
ZNF821
ONECUT2
KMT5C
ZNF516
ZNF708
SNCA
INSM1
FOXO6
AUTS2
PSIP1
E2F2
CHD7
ZNF397
HES6
MNX1
MYB
MAML3
SOX4
ZNF512
ZNF853
CBX2
ZNF704
APBB1
PAX5
TCF4
SETBP1
HOXD11
SALL1
FLI1
LHX2
SIX2
MYCL
TP73
SOX2
SOX21
DLX6
MYT1
L3MBTL4
ST18
HOXD9
HOXD8
MYT1L
FOXG1
ZNF93
ASCL1
HOXA9
HOXA4
SMAD9
HOXD13
ISL1
LMO2
NR4A2
HOXB8
CBFA2T3
ZEB2
HOXD10
TWIST1
NKX2-5
PAX6
POU4F1
GATA4
TBX1
MSC
LEF1
OVOL1
DLX2
IRF6
DLX1
RARG
SNAI2
AJUBA
LOXL2
DRAP1
FLNA
FHL2
TGFB1I1
CAVIN1
BASP1
TWIST2
DAB2
MITF
FGF2
GLI3
ENG
ZEB1
SMAD3
NOTCH2
EGFR
ELK3
EP300
CEBPB
STAT3
SGK1
TGFB1
FOSL1
GLIS3
BCL9L
MYBL1
RUNX1
TEAD2
RELB
RUNX2
HMGA2
WWTR1
YAP1
TEAD3
PTPN14
NFKB1
HMGA1
CCND1
PPARG
CDKN2B
NMI
LMNA
ETV4
FOS
CLU
JUP
KLF5
MUC1
EPAS1
GRHL1
BATF
RORC
TFAP2A
GATA3
TFAP2C
FOXP2
GRHL2
FGFR2
NKX2-1
HES1
OVOL2
FOXA1
ELF3
NOTCH3
SYK
EHF
BHLHE40

c

d

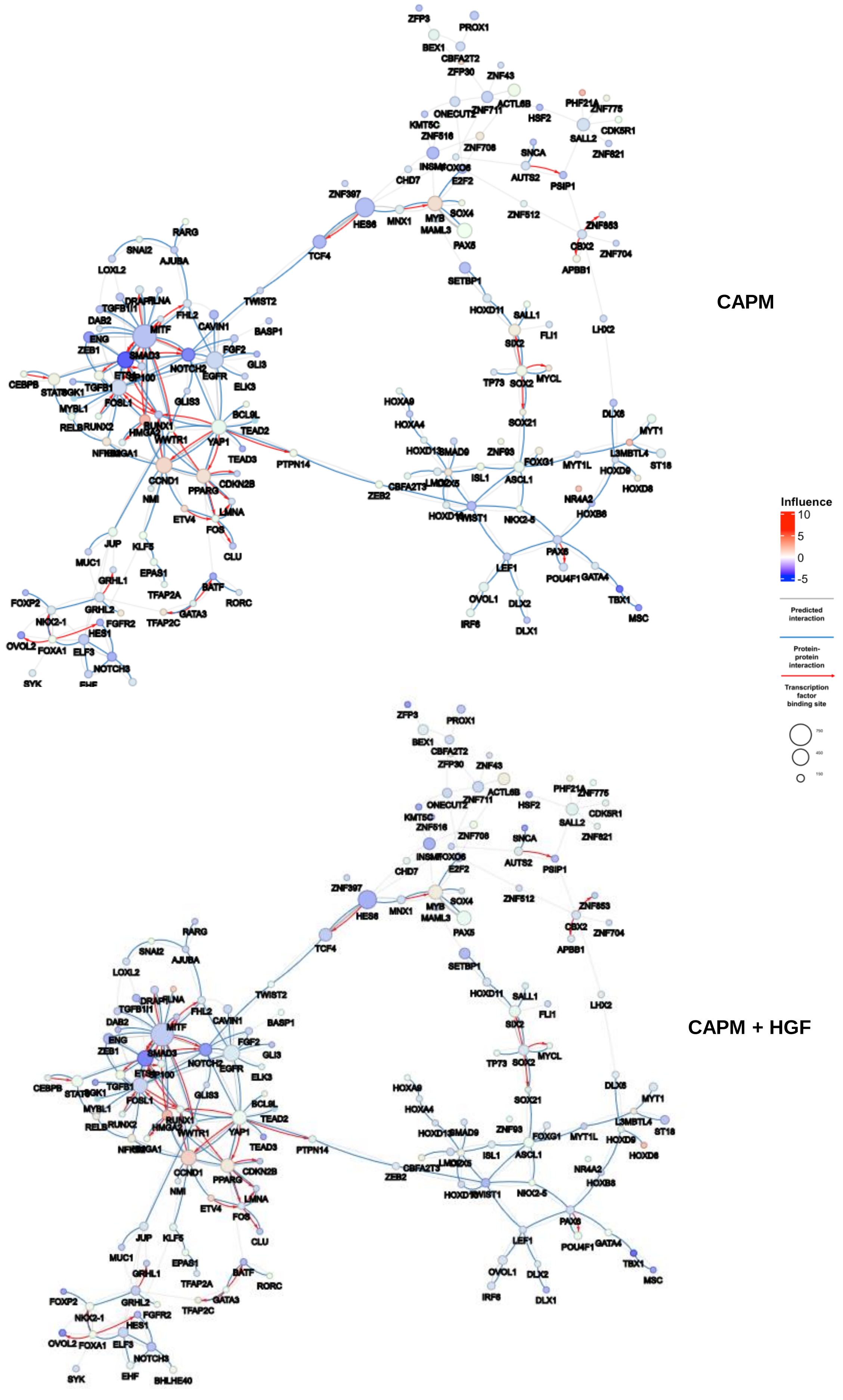
CAPM
CAPM + HGF
Influence
10
5
0
-5
Predicted interaction
Protein-protein interaction
Transcription factor binding site
750
450
150
ZFP3
PROX1
BEX1
CBFA2T2
ZFP30
ZNF43
ONECUT2
ZNF711
ACTL6B
HSF2
PHF21A
ZNF775
CDK5R1
SALL2
ZNF821
KMT5C
ZNF516
ZNF706
SNCA
AUTS2
PSIP1
INSM1
FOXO6
E2F2
CHD7
ZNF397
HES6
MNX1
MYB
MAML3
SOX4
PAX5
ZNF512
ZNF653
CBX2
ZNF704
APBB1
TCF4
SETBP1
HOXD11
SALL1
FLI1
LHX2
SIX2
TP73
SOX2
MYCL
SOX21
DLX6
MYT1
L3MBTL4
ST18
HOXA9
HOXA4
HOXD13
SMAD9
ZNF93
FOXG1
MYT1L
HOXD9
HOXD8
ISL1
ASCL1
CBFA2T3
LMO2
ZEB2
NR4A2
HOXB8
TWIST1
NKX2-5
PAX6
POU4F1
GATA4
TBX1
MSC
LEF1
OVOL1
DLX2
IRF8
DLX1
RARG
SNAI2
AJUBA
LOXL2
TWIST2
DRAP1
FLNA
TGFB1I1
FHL2
CAVIN1
BASP1
DAB2
MITF
ENG
ZEB1
SMAD3
NOTCH2
FGF2
EGFR
GLI3
ELK3
CEBPB
STAT6
SGK1
TGFB1
FOSL1
GLIS3
MYBL1
RELB
RUNX2
RUNX1
HMGA2
BCL6L
TEAD2
YAP1
WWTR1
TEAD3
PTPN14
NFIB
HMGA1
CCND1
PPARG
CDKN2B
NMI
LMNA
ETV4
FOS
CLU
JUP
KLF5
MUC1
EPAS1
GRHL1
TFAP2A
BATF
RORC
FOXP2
GRHL2
GATA3
NKX2-1
FGFR2
TFAP2C
HES1
OVOL2
FOXA1
ELF3
NOTCH3
SYK
EHF
BHLHE40

e

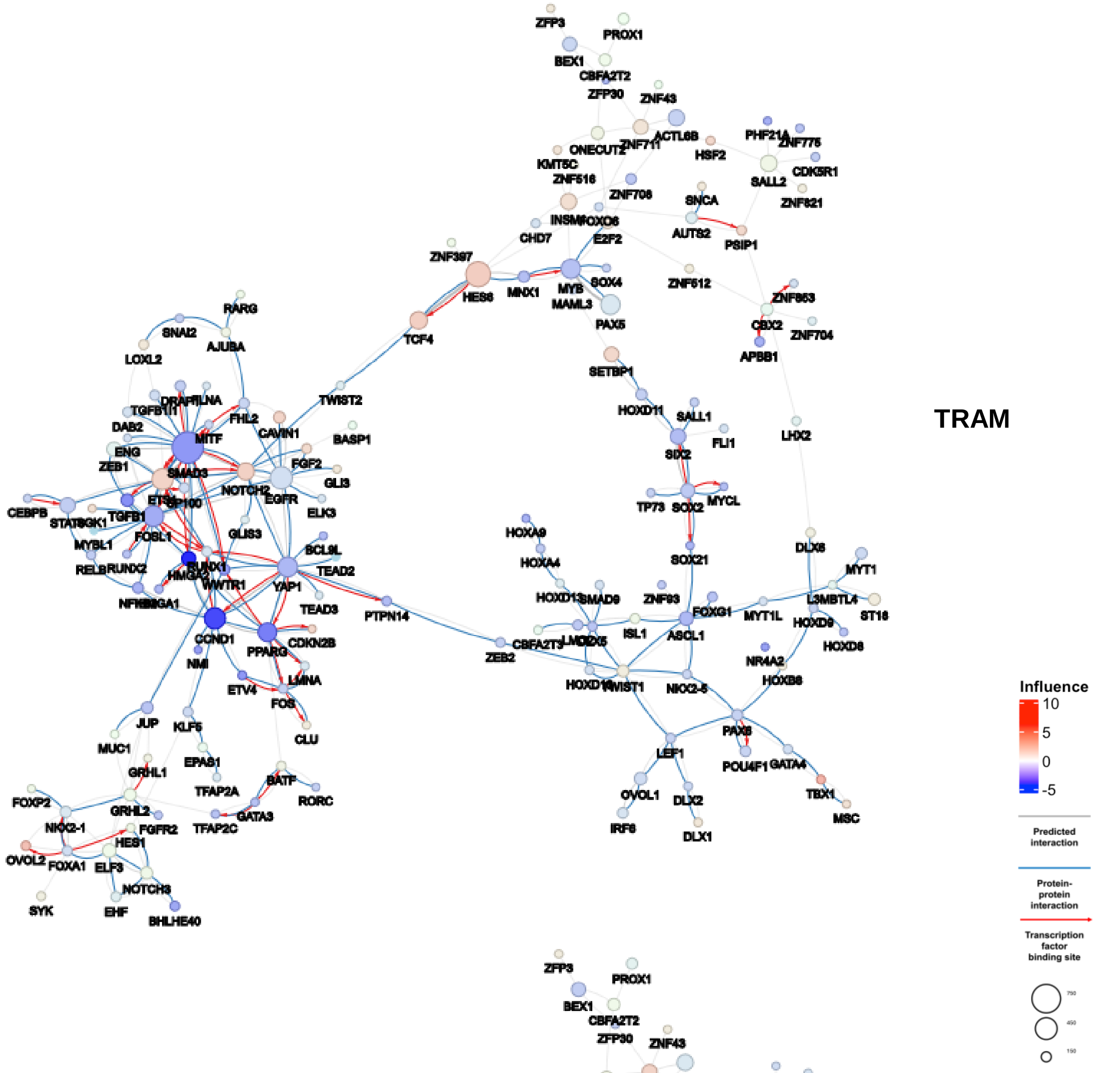
TRAM
Influence
10
5
0
-5
Predicted interaction
Protein-protein interaction
Transcription factor binding site
750
450
150

f

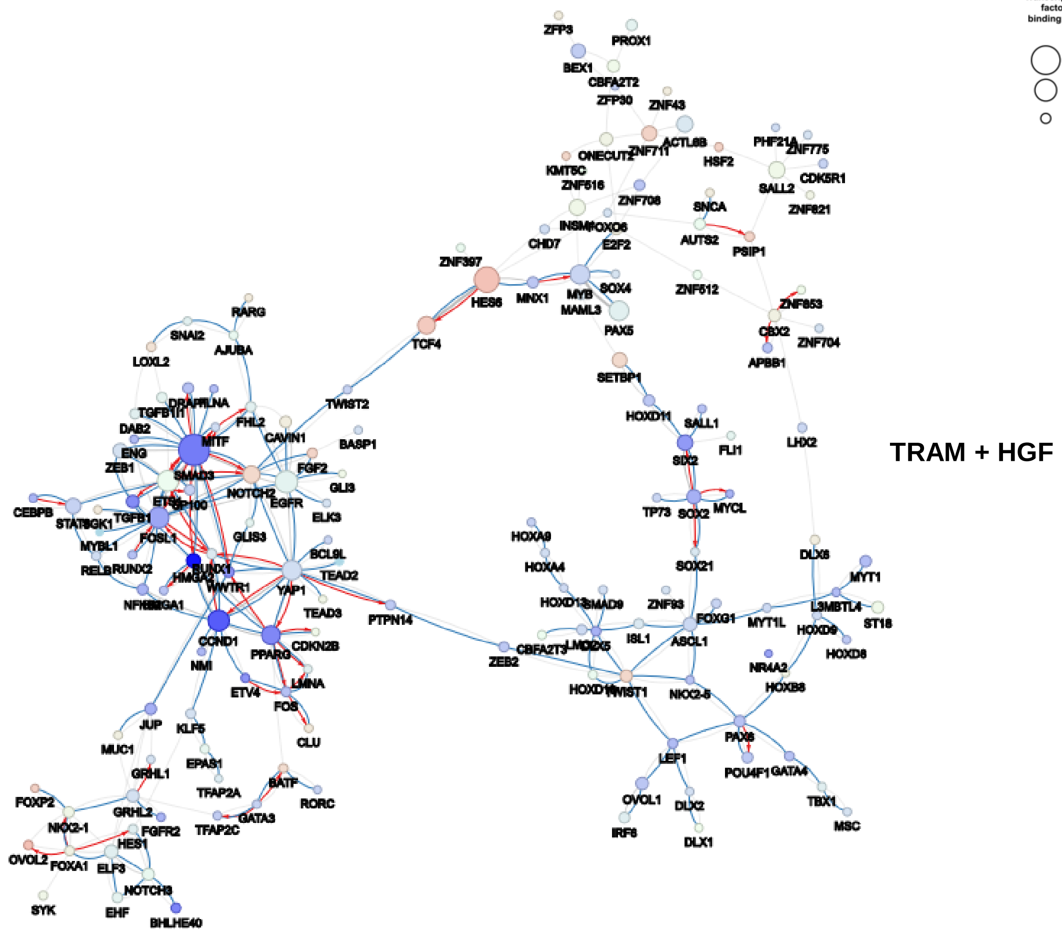
TRAM + HGF

**Supplementary Fig. S5 Regulatory network of METex14del in response to MET and MEK inhibitors.** Co-regulatory network showing the influence of TFs on METex14del cells without treatment in the absence (**a**) or presence (**b**) of HGF; or with treatment by capmatinib (CAPM) in the absence (**c**) or presence (**d**) of HGF; or with treatment by trametinib (TRAM) in the absence (**e**) or presence (**f**) of HGF. Circles represent DIRs and the radius of the circle is proportional to the number of target genes regulated by the DIR. Co-regulatory interactions between DIRs are indicated: protein-protein interactions with published evidence (blue lines), transcriptional regulation interactions with published evidence (red lines), and interactions defined only by the h-LICORN algorithm (gray lines).
