## Supplemental Tables for "Transcriptional program-based deciphering of the MET exon 14 skipping regulation network"

**Supplementary Table. S1** Primers sequences used for qPCR

| **Target** | **Forward Primer** | **Reverse Primer** |
|---|---|---|
| B2m | GCTGGCGGGCATTCCTGAAG | TGCTGGATGACGTGAGTAAACCTG |
| ETS1 | AGTCGATCTGGAGCTTTTCCC | AGTCCCGAACATGGGTTTCT |
| FOSL1 | GGCGGAGACTGACAAACTGG | TTGGCTCCTTCCGGGATTTT |
| SMAD3 | GATCCCACCAGGATGCAACC | CTTTGACGAAGCTCATGCGG |
| HMGA2 | TTCATGCCATATGCCCCATCC | TTCTTGGAGCTGGTTCTTGGT |
| CCND1 | TGCTGCTATTGGAGGATCAGTT | TGTGAAACAGCAACCTTTTTGGA |
| RUNX1 | CCCGGAGGGAAACTGTGAAT | CGAGTTGCATCCCTCCTCTC |
| MMP1 | GTTCAGGGACAGAATGTGCTA | CTGCAGTTGAACCAGCTATTAG |
| VIM | ACGAAACTTCTCAGCATCACG | TTGCGCTCCTGAAAAACTGC |
| KRTAP2-3 | CTGGCCCTCTCTGACAATCTT | TGATGAGTCAGTGGGACAGAG |
| NGEF | CCCTGATGGGCTGGATGTAA | CGATACAAGATGCAGTGACCA |
| NOG | ACCAGTTCCACCACCCTCTA | ACTGGGACCGTATATACACACA |
| SERPINE2 | AGAAACCATGCAAAGCAACG | ACTTGCATCGAGTCTGTTCCT |
| LETM2 | CCATTGTCACCCCACTCAACT | TCCAAGCCATAGGAGACAGGT |
| SH2D5 | ATCCCTCCAGCCAGCATAGA | GTGGCGCAGATTGGAACAAA |
| TSPAN5 | GTATGAAAGTCGTTGCGCCG | TCCAAGGTGTGGTGAAGAGC |
| G0S2 | CCACAAGCATCCACCAAAGG | ATCCTTCCTCCCTAGTGCAA |
| ADAM19 | TCTAGACCTGTCCAAGGGGC | GGCCAGACATGCTTCTTCAG |
| RAC2 | CGGTGAAATACCTGGAGTGCT | TGCCATCAGGCTTTGGGTG |
| CAVIN3 | GGAGCGAATCCTACATCCACC | TTATTGATGGTGAGCGCAAGC |
| COL13A1 | TTGTGTCCTGGTGCCAAAGG | TCTTAGACACACCTAGCTTAGCA |
| ARHGAP22 | CTTTTGGGACCCCACACACT | GGCTAAGCGCTACTCTTGGT |
| THBD | CACACAGGGTCTGCAAGGTC | AGCAAGGATTTTGCCTGTGTT |
| ABCA1 | CCAAAGAGCCATGTGTCATGT | ACATTCTAAGGGGGATGCAACA |
| DCBLD2 | TGGGAGAGAGAGTTCGCATCA | TTCCACTCATGAACAGCAATGTG |
| PTX3 | CATGCCAGTTGGGAAGGTCTG | CTGAAAGCACCATGGCATAAAGT |
| CHGB | CTACGGTGAGGAAGGAGCCC | AAAATGGCTGCTCTTCCACTG |
| SERPINA1 | CTCCCCTCTTCATGGGAAAAGT | GGACCAGCTCAACCCTTCTTT |

**Supplementary Table. S2** Influence of key differentially acting regulators (METex14del+HGF vs. resting conditions)

| DIR | MeanGroupInfluence | MeanRestInfluence | AdjPval | type |
|---|---|---|---|---|
| HMGA2 | 4,5508 | -1,8105 | 4.17e-04 | Over |
| ETS1 | 3,8761 | -1,4845 | 2.84e-03 | Over |
| SOX9 | 2,8294 | -1,2940 | 8.17e-03 | Over |
| CCND1 | 2,4282 | -0,9955 | 4.81e-02 | Over |
| FOSL1 | 2,3546 | -1,0999 | 4.19e-02 | Over |
| HOXD4 | 2,3460 | -0,8551 | 3.32e-02 | Over |
| RUNX1 | 2,1534 | -0,8914 | 9.16e-02 | Over |
| NR4A1 | 2,1055 | -0,5357 | 7.66e-03 | Over |
| SMAD3 | 1,9699 | -0,8205 | 4.75e-02 | Over |
| TGFB1 | 1,9444 | -0,8192 | 1.21e-02 | Over |
| SALL1 | 1,9127 | -0,6528 | 2.55e-02 | Over |
| HOXD3 | 1,8624 | -0,8141 | 2.02e-02 | Over |
| MSC | 1,8353 | -0,6275 | 7.09e-03 | Over |
| ZEB1 | 1,7396 | -0,7930 | 2.78e-02 | Over |
| WWTR1 | 1,6920 | -0,7395 | 6.83e-03 | Over |
| HOXD9 | 1,6884 | -0,8022 | 1.91e-02 | Over |
| MYB | 1,6128 | -0,6217 | 1.46e-02 | Over |
| HOXD13 | 1,5416 | -0,6198 | 4.75e-02 | Over |
| HOXA4 | 1,4245 | -0,8688 | 1.11e-02 | Over |
| MNX1 | 1,3437 | -0,5172 | 4.19e-02 | Over |
| ZNF43 | 1,3317 | -0,5225 | 3.38e-02 | Over |
| PAX6 | 1,3268 | -0,5624 | 2.54e-02 | Over |
| MYT1 | 1,3089 | -0,5441 | 3.68e-02 | Over |
| HOXC10 | 1,2486 | -0,4473 | 3.32e-02 | Over |
| BNC1 | 1,2148 | -0,5059 | 1.91e-02 | Over |
| ZNF257 | 1,1861 | -0,5422 | 9.98e-03 | Over |
| ELK3 | 1,1728 | -0,4717 | 7.31e-01 | Over |
| MECOM | 1,1120 | -0,4340 | 1.16e-02 | Over |
| S100A8 | 0,9706 | -0,5784 | 2.13e-02 | Over |
| GLI3 | -0,8140 | 0,3991 | 2.76e-02 | Under |
| CRABP2 | -0,9510 | 0,3962 | 1.46e-02 | Under |
| RARG | -1,0230 | 0,3897 | 3.94e-02 | Under |
| MAGEA2 | -1,1677 | 0,3997 | 8.12e-03 | Under |
| IRF9 | -1,2384 | 0,5029 | 9.98e-03 | Under |
| MAGEA2B | -1,3764 | 0,6316 | 3.66e-03 | Under |
| SP100 | -1,4076 | 0,5079 | 3.99e-02 | Under |
| POU6F2 | -1,4444 | 0,5135 | 7.06e-03 | Under |
| HSF2 | -1,4562 | 0,6733 | 1.25e-02 | Under |
| TEAD3 | -1,4950 | 0,4976 | 7.06e-03 | Under |
| FOXF2 | -1,5980 | 0,7195 | 6.39e-03 | Under |
| SIX3 | -1,6073 | 0,8518 | 6.56e-03 | Under |
| FLI1 | -1,6801 | 0,5893 | 4.60e-02 | Under |
| IFI27 | -1,7500 | 0,7008 | 2.84e-03 | Under |
| GRHL2 | -1,8636 | 0,7035 | 2.62e-02 | Under |
| IRF6 | -1,9829 | 0,9480 | 1.46e-02 | Under |
| NOTCH3 | -2,0048 | 0,6824 | 4.75e-02 | Under |
| SP110 | -2,0192 | 0,8257 | 2.02e-02 | Under |
| HOXD10 | -2,0514 | 0,7144 | 1.04e-02 | Under |
| DZIP1 | -2,0892 | 0,7596 | 2.44e-02 | Under |
| MAGEA1 | -2,1285 | 1,1628 | 4.18e-03 | Under |
| SNAI2 | -2,2596 | 0,9031 | 6.83e-03 | Under |
| AJUBA | -2,3348 | 0,8951 | 7.06e-03 | Under |
| TFAP2A | -2,3414 | 0,8126 | 5.20e-03 | Under |
| NKX2-1 | -2,7241 | 0,9467 | 4.25e-04 | Under |
| FOXA1 | -2,7561 | 1,0003 | 7.65e-04 | Under |
| TRIM29 | -2,7605 | 0,9482 | 4.17e-04 | Under |
| EPAS1 | -2,8077 | 1,1955 | 6.39e-03 | Under |
| ELF3 | -2,9620 | 1,2628 | 1.46e-02 | Under |
| TWIST1 | -2,9774 | 1,1388 | 2.84e-03 | Under |
| MYO6 | -3,0274 | 0,8719 | 1.50e-03 | Under |
| LOXL2 | -3,5311 | 1,4055 | 5.28e-03 | Under |

**Supplementary Table. S3** Genes targeted by ETS1, FOSL1 and SMAD3

| TF | Regulation | Target Genes |
|---|---|---|
| ETS1 | activated | CFLAR,MAP3K14,IFFO1,MRC2,ANLN,CD74,SAMD4A,RIPOR1,LIMA1,DCBLD2,SLC9A7,IFI35,ACSL4,NEDD4,STK10,CD59,FXYD5,ITGA6,CCDC80,GADD45B,LGALS1,APOL1,MYH9,NALCN,NDRG1,SYDE1,CDK6,NAMPT,SERPINE1,DNMBP,DKK1,LGALS3BP,DHX58,SMURF2,PMP22,RAB34,CPZ,KLHL5,WNT5B,SH2B3,GNAI2,GNB4,GBE1,MAPKAPK3,CLIP4,GADD45A,UAP1,GBP1,GDA,OGFRL1,IFIT3,CD274,TNFRSF10B,BICC1,RASSF8,ACOT9,SDC4,TRIP10,ARMCX1,FLNC,CALU,ARHGAP22,MAP7D3,CRACR2A,SH3BGRL,RFTN1,XAF1,STARD13,RRAS2,NGF,IL6ST,IL15RA,SLC43A3,RAB11FIP5,DYSF,FAM129A,DOCK10,DDX60,ARHGAP29,IFI44,AOX1,HERC6,FGF5,FRMD6,PCSK6,IFITM3,COL6A1,EMP3,EFHD2,SH3BGRL3,EVA1B,CYR61,ABL2,ACKR3,OSBPL10,PTPRG,COL8A1,NCEH1,LHFPL2,RASA1,PPP1R18,ARHGAP18,SH3KBP1,MSN,SLC25A37,UGCG,RGS10,TAGLN,CDC42EP2,PRSS23,ME3,NMT2,UBASH3B,ROBO3,RHOC,MAP3K7CL,UBE2L6,TMEM171,MMP14,ST3GAL2,SLC34A2,TNFRSF14,WDR66,GALNT14,ZYX,ITGA5,NEXN,CDC42EP3,IFI16,PTX3,ANXA5,EDIL3,ANKRD33B,GPX8,CGAS,DLC1,PDP1,ABCA1,LETM2,GPR176,RAB8B,SERPINB8,SLFN5,CERCAM,TPM4,PXK,FILIP1L,TSPAN5,PXDC1,CLIC4,MT1E,TM4SF1,IFFO2,NINJ2,SLFN11,TNFRSF10D,RIN1,NRIP3,C11orf68,SPHK1,OXTR,F2R,BACE2,KIRREL1,PRR16,IFITM2,KCNQ5,RASA3,SEPT10,TRABD2A,S100A3,SH2D5,HRH1,ANXA6,PDCD1LG2,NMB,LPAR1,ECI2,PLXNB3,TAP2,MICB,SAMD9,GPSM3,TAX1BP3,APOBEC3G,PNMA2,TMEM35B,LYN,LIX1L,PRSS2,KRTAP2-3 |
| ETS1 | repressed | SEMA3F,TMEM176A,KLHL13,ICA1,ABCC8,TTC22,USH1C,SCIN,CACNA2D2,IL20RA,CNTN1,SYT13,PLEKHB1,NLRP2,GRAMD1B,CDH1,FSTL4,ENTPD2,SYNE2,KCNH2,GYG2,PSD,SEZ6,ATP2C2,ABCA7,STAG3,REEP1,ATP11A,DAPP1,ST6GALNAC1,LLGL2,ATP2A3,MAP2,IGSF9,TTC39A,CEACAM6,CHRD,NRCAM,SLC7A8,HOOK2,BAMBI,DSP,CECR2,TRIM9,BMP7,NOL4,SYP,ELMO3,MMP15,TOX3,BMF,SCG3,FZD3,NCALD,TLE6,CAPS,LSR,PON3,CHN2,PTPRZ1,GLCCI1,TSPAN13,TMEM176B,RUNDC3A,MAP2K6,CPE,DDX25,DTX4,CALCA,RASAL1,ADGRD1,ENPP5,HMGCS1,PDE4D,HGD,WNT5A,IGFBP2,RGS2,SIPA1L2,SGIP1,PTK2B,EPHX2,PRXL2A,ATP7B,POF1B,RAB17,HSPA2,TSPAN8,FBXL16,DLL4,SYT5,APOE,GADD45G,SMPDL3B,RAB11FIP4,GCH1,DDC,PATJ,SYT4,PIK3C2B,LARGE1,FAM83F,NTS,UNC79,CMPK2,BICDL1,CGA,PRRG4,GLS2,ARHGEF4,CIB2,SULF1,FXYD6,GCHFR,B4GALNT3,FAM222A,TTC6,AK7,CCDC33,TMC6,RNF165,SIK1,CNKSR1,RGS16,MINDY1,VASH2,SLC27A3,STXBP5L,N4BP3,CPLX2,VWDE,ADGRL3,LYPD6B,AKR1C2,MFSD6,RNF144A,TMEM163,C4orf19,SORBS2,ELMO1,MPV17L,TSPAN33,STC1,BTG2,CELF3,KALRN,PDE9A,SPTBN4,MPZL3,CHRNB2,ELAVL4,WNT4,KCNF1,FAM84A,VSNL1,SGPP2,EIF4E3,TMEM169,PPM1L,CDS1,RBM47,HPGD,CMBL,KCNK5,DEFB1,ALDH1A1,WNK2,AQP3,PHYHIPL,NDRG2,PPFIBP2,RIC3,CRABP1,SEC11C,PLEKHA7,MS4A8,MYO5B,TMEM145,SPINT2,HID1,RAB26,DEGS2,MFSD2A,NPNT,ENHO,PARM1,LRRC45,GTSF1,GPR27,NRTN,TMEM37,KCNK3,RND1,AGR3,MFSD4A,ABO,PCSK1,LPL,CALCB,CRYBG2,GPX2,PRR15,TCIM,RIMKLA,SRRM3,GLDC,MKRN3,NRXN1,IDH2,PCP4,BEGAIN,RIPPLY3,CCSER1,MAP7D2,PLA2G6,KCND2,KCNH7,OSBP2,RPS27L,C12orf56,TMEM105,MARC1,CYP4F3,CARMIL3,AKR1C1,RINL,SBK1,TMEM198,ZDHHC11,RASSF10,SEMA4A,EVL,KCNMB2,NUP62CL,DLL1,PPP1R14C,SMOC1,SFMBT2,IQANK1,RBM20,CLDN9,UBE2QL1,CKMT1A,CKMT1B,CASTOR3,SELENOP,MYO15B,CBSL,SRCIN1 |
| FOSL1 | activated | CYP26B1,CFLAR,COPZ2,ITGA3,MAP3K14,TNFRSF12A,IL32,FHL1,BIRC3,TMSB10,CAPG,ADAMTS6,CYBA,CTSA,SLC9A7,CA12,FGFR1,SYNJ2,EPB41L2,SLC4A4,CD82,PPP1R15A,ITGA6,CARD10,LGALS1,TSPO,PLTP,CD40,TIMP1,KLHL4,NDRG1,CDK6,CAV2,DKK1,RAB34,SLC15A3,SH2B3,ADTRP,HBEGF,GNAI2,GBE1,GADD45A,F3,NRP2,TNFAIP3,CTNNAL1,IFI27L2,PPP1R3C,PYROXD2,CD274,PDLIM2,NPPB,INHBA,SERPINB6,ATXN1,CPA4,FLNC,TWSG1,CD68,KLK10,MAP7D3,ATP8B3,C19orf66,SLC35D2,AKAP12,RFTN1,PDLIM4,XAF1,EDEM1,FST,IL15RA,CLDN10,OASL,DYSF,BIN1,IER3,ARHGAP29,CALHM2,CDK15,HERC5,CDH11,IFITM3,COL6A2,ADAMTS10,EVA1B,TM4SF19,SLC25A37,RGS20,ITPRIP,ADAM12,RGS10,SERPING1,JPH2,LYPD1,GJA1,RBMS1,UBASH3B,ROBO3,ANGPT1,PITPNC1,SH3RF2,MMP14,AFAP1L1,GALNT14,NBL1,DMTN,LY6K,FBXO27,FBLIM1,MEGF6,FSTL1,NUAK2,ZC3H12A,TMEM144,ERAP2,CSF2,IL31RA,HTRA1,GPR176,LARP6,RRAD,NNMT,PXK,RAB31,TSPAN5,MT1E,TM4SF1,IFFO2,KISS1,CAVIN3,PRNP,CALB2,SPTLC3,EFEMP2,ABLIM3,SAA1,TNFRSF10D,C11orf68,TUBB6,SPHK1,FJX1,MXRA7,PAQR7,COL18A1,CCBE1,UPP1,KIRREL1,SMIM10,FMNL1,EVI2B,KRT5,S100A3,PLEKHG4,SLC6A9,MMP1,PDLIM7,SLC22A4,COL13A1,TPM2,PLXNB3,TGM2,MICB,DIO2,ARHGEF28,FADS3,PSG1,C2orf74,PSMB9,APOBEC3C,TMEM158,LYN,NBPF14,F8A3,KRTAP2-3 |
| FOSL1 | repressed | GCLC,SEMA3F,TMEM98,USH1C,ARHGAP44,KDM5D,IL20RA,CNTN1,SYT13,NRXN3,SH3YL1,GPM6B,XK,ATP9A,GYG2,CDK14,LIMCH1,EYA2,ERBB3,KIF26A,REEP1,SPTB,ST6GALNAC2,MGAT4A,FRY,ANO8,ARHGEF10L,PLXNA2,GPC4,MAP2,LAMP3,NEBL,TNS1,SCGN,NID2,PHACTR3,SULT2B1,SLC4A11,CHGB,YPEL3,SEMA6A,ABLIM1,PALMD,CECR2,SEPT3,SYNGR1,CHGA,RIMS4,NOL4,FGF9,PARD6A,HOMER2,SCG3,FZD3,STMN2,NCALD,MCM4,TLE6,CADM4,TSPAN12,CHN2,COBL,TMEM176B,SPOCK2,SH3PXD2A,DNAJC12,MTMR4,ENO3,DDX25,B3GAT1,LIN7A,ADGRD1,TBC1D30,EYA4,HMGCS1,RBP1,FGF12,FRMD4B,RTKN,IGFBP2,IGFBP5,EFHD1,CHST10,KIAA1324,RHOU,C1orf21,BMP8B,PIK3R3,MAN1C1,SGIP1,TMPO,SOCS2,CNTFR,MAGEA10,FLRT3,AMOT,HSPA2,AIF1L,FGD3,TSPAN8,TNFRSF19,CGNL1,TNNI3,KIF1A,NAPSA,C19orf57,SERPINF1,PIK3C2B,NTS,VAV3,CMPK2,GRP,EPHA7,MAP7,USP44,GPNMB,RALGPS1,TMOD1,IGFBPL1,FXYD6,SEMA6D,PARP9,MARCH9,SLAIN1,STON2,MISP3,HUNK,TRPM2,SLC44A3,CELSR2,CGN,STXBP5L,SCD5,SSBP2,DCDC2,LGSN,SHROOM2,CACNA1B,INA,GSTO1,GLB1L2,DOC2A,TMEM45B,LPCAT1,GDPD1,JPH3,TSPAN7,PCDH1,CACNA1D,CACNA2D3,DYRK1A,TNFRSF14,RADIL,CACHD1,EPB41,TMPRSS3,CHRNB2,MEIOB,BEND5,ELAVL4,ATXN7L2,TRIM58,C1orf115,DAPL1,TMEM169,CADPS,YEATS2,ABLIM2,LMBRD2,ADGRV1,TERT,GRIK2,TMEM74,TMEM184A,BAALC,CLDN3,NDRG2,LARGE2,DCHS1,CRABP1,RAB26,SPINDOC,KIAA1586,HR,NPNT,NSG1,ENHO,PARM1,COL22A1,SDC2,NRG4,LINGO1,RNF150,LONRF2,GTSF1,GPR37,GPR27,KCNK3,HOPX,ID4,CSRP2,PCCA,PCSK1,CRYBG2,FAM89B,RIMKLA,CA8,RAB39A,PACS2,MKRN3,NRXN1,LYNX1,LRRC75A,IDH2,CEP97,CADM1,PCP4,ABAT,RIPPLY3,UTY,CTAG1B,USP18,MANEAL,METTL7A,IFNLR1,BACE1,SLIT1,KCNJ11,SBK1,NKAIN2,TMEM198,KLRG2,FAM111B,TCEAL3,SULF2,SLC25A29,KCNMB2,ATL1,KLHL9,FAM169A,EFCAB2,COL5A2,RGL3,DENND1C,HBA1,LBH,COLCA2,FAM19A5,SFTA3,PNMA6A,CASTOR3,TMEM150C,SELENOP,MAGIX,C2orf15,CBSL,NEFL |
| SMAD3 | activated | CYP26B1,SLC7A2,COPZ2,LAMP2,MRC2,PLAUR,ANLN,CLDN11,MVP,GPRC5A,MAMLD1,CYP24A1,FHL1,SNAPC1,GCLM,VIM,FAS,CD44,TMSB10,VCL,RAB27B,TNC,ADAMTS6,DKK3,LIMA1,CYBA,DCBLD2,BCAT1,CASP8,FAM107B,TNFRSF1A,NAV3,IFI35,ACSL4,FSTL3,LMCD1,ACTN1,SYNJ2,LXN,CD59,PTHLH,PXN,ICAM1,LAMB1,MAP3K20,TGFB2,NRP1,GADD45B,PLEK2,RIN3,PROCR,CD40,CTSZ,ASB9,TIMP1,KLHL4,COTL1,FAH,PLAT,TNFRSF10A,PLIN3,GSDME,CAV1,MET,PTGR1,ACTA2,SFXN3,MAP3K8,ALDH3A1,ICAM2,SMURF2,CPZ,KLHL5,CTSC,EHD1,SLC15A3,TNS2,CAP2,CRYBG1,ADGRG6,HBEGF,PDE4D,STC2,COL7A1,GNAI2,PLSCR4,IL1A,STEAP3,CLIP4,EFEMP1,MLPH,QSOX1,NID1,UAP1,GBP3,GBP1,MUC5B,SPP1,IFI27L2,LTBP2,IFIT3,PPP1R3C,ARAP3,TGFBI,TNFRSF10B,TRIM6,CSTA,PLAU,CHST3,RASSF8,ACOT9,OPTN,SDC4,SERPINB6,NQO2,ATXN1,EREG,AHNAK,MT2A,TNFSF9,TRIP10,EVI2A,SLC10A3,ARMCX1,ZC4H2,APOL3,RAC2,LIF,KRT17,CPA4,CALU,LOXL1,PALLD,C19orf66,MPP1,ULBP2,GFPT2,TRIM21,RAMP1,MATN2,RIN2,STARD13,EPHB2,MICAL2,IL6ST,FST,RTL8C,SLC37A2,CTSL,ADAM19,OASL,NT5E,RAB11FIP5,FAM129A,SERPINE2,FLNB,PHF11,TNS3,FAM129B,SLC31A2,DDX60,CASP1,THBS1,BCAR3,IFI44,MYOF,STAMBPL1,DUSP5,CALHM2,GPAT3,CPNE8,PRICKLE1,AMIGO2,PHLDA1,FRMD6,KIFC3,CMTM3,CDH11,MEAK7,IGFBP4,FKBP10,COL6A1,COL6A2,EPHA2,EFHD2,SH3BGRL3,CYR61,ABL2,PHLDB2,NCEH1,TM4SF19,OCIAD2,PLAC8,OSMR,PLK2,LHFPL2,ARHGAP26,ARHGAP18,SH3KBP1,SLC16A2,UGCG,ANKRD1,GSTO1,PTPRJ,TAGLN,CDC42EP2,ITGB1,LYPD1,VEGFC,IL18,CRIM1,TDO2,GJA1,PLOD2,CMTM7,ANKRD29,ANGPT1,PRKCA,GRAMD2B,RHOC,FZD7,KCNMA1,SH3RF2,CD109,FBXO32,RNF207,RHBDL2,CDA,ZYX,SHC1,ITGA5,DHRS3,NEXN,SNX7,GBP2,CAPN2,CDC42EP3,ANTXR2,TGFBR2,IFI16,PTX3,MELTF,SPRY1,ANXA5,IL15,ITGA2,EDIL3,F2RL1,GPX8,CGAS,IL31RA,GALNT10,CTSB,CTHRC1,ABCA1,LETM2,ZCCHC24,BEND7,JCAD,UBTD1,HACD1,SPRED1,FBN1,LARP6,CLMP,SERPINB7,SERPINB8,NNMT,SLFN5,ANPEP,TMEM92,TPM4,AXL,IGFBP6,LTBP3,MLKL,ADAM9,IL7R,PXDC1,PCSK9,PLEKHA2,SERPINB9,EMB,CAVIN3,CHST11,NINJ2,PRNP,SYNPO,EPHX4,LAMB2,SPTLC3,GNG12,EFEMP2,GLRX,SNCG,MMRN2,HEG1,RIN1,POLD4,TUBB6,CD151,ODF3B,PLEC,RNF212,THBD,CPNE7,CDH4,MYADM,OXTR,F2R,PHLDA2,LDOC1,EXT1,MXRA7,EPGN,ANXA2,COL18A1,GPR1,NOG,LHFPL6,GPR39,ARSI,OAF,ALDH1A3,SOCS3,PRR16,SEMA4B,SIGIRR,TNFAIP2,TLCD2,AHNAK2,ZFP36L1,KCNQ5,EVI2B,TRABD2A,COL4A1,COL4A5,PLEKHG4,AFAP1,MME,FAM3C,SERPINA1,PDCD1LG2,PARVA,S100A10,ITGBL1,RUSC2,SAMD9,KRT6A,S1PR3,CLIC1,FADS3,GPX1,TNFSF12,APOBEC3G,AKAP2,PSG4,H2AFJ,FMN1,LUZP6,CCL5,PRAG1,PRSS2,HLA-E |

| | | |
|---|---|---|
| SMAD3 | repressed | TMEM176A,LIG3,SPPL2B,ABCC8,PRSS21,PROM1,CACNA2D2,SYT7,ALOX5,NLRP2,PRKCH,CDH1,GPM6B,MCF2L2,SYNE2,FAM168A,KCNH2,PHF21B,PTPRU,CDH3,SEZ6,EYA2,STAG3,SPTB,ASNS,ST6GALNAC2,LNX1,LLGL2,ADAM11,CACNG4,WDR62,RAP1GAP,PPP1R12B,BRINP1,CEACAM1,PAFAH1B3,COL4A4,ERC1,KAT6A,FAM234B,SLCO1A2,NKAIN1,CEACAM6,SULT2B1,SLC4A11,NRCAM,COL9A3,MSH2,CDC7,ARVCF,PRODH,NEFH,KCTD17,COCH,TRIM9,TMED8,DAAM1,CHGA,EFS,CCM2L,CELF4,CBLN1,NDRG4,ESRP2,HSDL1,TOX3,QPRT,XYLT1,AP3B2,NCALD,KLC3,TJP3,MYH14,LIG1,CAPS,FAM83E,RAB3A,ITGB8,PTPRZ1,GLCCI1,CEP41,RLN2,RGP1,SPOCK2,FBXL15,RASD1,KIAA1211,MAPK10,CPE,CRYAB,NRXN2,TRIM3,MADD,NAA40,IL23A,RAD51AP1,SCNN1A,TBC1D30,MDM1,FAM184A,EYA4,MDFI,PODXL2,UPK1B,PEX5L,RTKN,PASK,EPHA4,MARK1,AGMAT,KIF21B,SIPA1L2,ARTN,PIK3R3,BSPRY,GRIA2,DNAJC15,HPCA,PRXL2A,ENKD1,FAM210B,ATP8A1,POF1B,MOCS1,CDKN1A,NRN1,SH2D3A,FAM110A,ST3GAL3,KLC1,AIF1L,CTAG2,FGD3,VIL1,TNFRSF19,RASL11B,GAL3ST1,DLL4,PIMREG,EPB41L4A,APOE,APOC1,GADD45G,GTPBP3,UNC13A,ASS1,LRP3,EPS8L1,ACAP3,GCH1,DDC,RAB25,SYT4,CHRM3,DCLK1,FAM83F,UNC79,TTC9,GSTM3,RERG,TUT4,DSC2,FHOD3,MSI1,BICDL1,DTX1,CGA,MAP7,ALDH1L2,RAPGEF5,CTSV,RNF38,MDC1,SULF1,SORL1,TMPRSS4,SEMA6D,FNBP1L,TACC2,FRAS1,SHROOM3,KIF21A,FGD4,TTC6,CDH24,NOVA1,AK7,DNAJA4,CCDC33,RNF165,GRB7,LMTK3,SLC44A3,RGS16,CGN,HCN3,PARP1,TMEM108,RUBCN,MUC4,SLC10A4,N4BP3,CPLX2,SDK1,ATP6V1B2,INA,ST14,MPZL2,ADGRL3,AKR1C2,RNF144A,TMEM45B,SERP2,ANKRD22,RAB3C,NRSN1,ACOXL,SYCP2L,BMP6,RETREG1,CCSAP,SRSF12,FBXO43,MPV17L,SSBP3,CACNA1D,TNFRSF14,RADIL,BTG2,CELF3,FNDC5,SLC37A1,GATD3A,SPTBN4,BDH1,TBC1D24,BICDL2,SLC25A45,ZSWIM5,WNT4,FLVCR1,VSNL1,SLC16A14,TMEM169,IFT122,PLXNB1,BSN,SLC29A4,TMEM184A,CLDN3,WNK2,AQP3,PHYHIPL,FAAH2,TC2N,MOAP1,NELL1,SPINT1,GABRB3,SEC11C,CATSPER2,KATNAL2,ACAA2,PRRT2,CORO6,RCOR2,DEGS2,MFSD2A,FAM110B,COL22A1,CCDC8,NRG4,KCNAB3,DNAJC18,PPM1D,MTSS1,GSTA4,HS6ST2,MAP6,GPHN,C2CD2L,IL17D,ZNF738,CES4A,CES3,NBEA,AGR3,CCDC106,KLHL15,CNTNAP2,MFSD4A,SLC29A2,CNIH2,PDIK1L,UCP2,CALCB,DOK7,HSD11B2,RIMS2,PRR15,MYO1D,TCIM,POLE,TOP3A,PPFIA3,ERICH5,SRRM3,GLDC,TMEM52,NSUN7,ANKRD18A,MTURN,MAP6D1,FAM83H,HIST3H2A,C6orf223,CLDN7,RTKN2,TMEM30B,EPHB3,SPNS2,PCP4,ABAT,NEB,PRR36,EPHA10,RIPK4,MACC1,FAM3B,UTY,MAP7D2,PLA2G6,KCND2,KRBA2,RPS27L,C12orf56,CYP4F3,CARMIL3,BCAM,MAGEH1,FANCA,RALGAPA2,FBLL1,FAM221A,KLRG2,CLDN4,RASSF10,GREB1,RYR1,SULT1A1,HSH2D,MAGEA6,INKA2,SVIP,TEX45,SFMBT2,CSAG1,FAM229B,RBM20,PTPN20,HSD17B8,MROH6,PDE7A,SAP25,ADGRG1,DENND1C,RTL8B,RNF208,KLHL23,MAGEA12,EML6,UBE2QL1,TSTD1,ATAD3C,FAM19A5,MAGEA3,FNDC10,KIAA0040,PNMA6A,CKMT1B,KRBOX1,ARHGAP8,ARHGDIG,MC1R,XKR7,EPPK1,MYO15B,SMIM22,CTAG1A,CD24,SIK1B,ERVMER34-1 |
| ETS1-FOSL1 | activated | CFLAR,MAP3K14,SLC9A7,ITGA6,LGALS1,NDRG1,CDK6,DKK1,RAB34,SH2B3,GNAI2,GBE1,GADD45A,CD274,FLNC,MAP7D3,RFTN1,XAF1,IL15RA,DYSF,ARHGAP29,IFITM3,EVA1B,SLC25A37,RGS10,UBASH3B,ROBO3,MMP14,GALNT14,GPR176,PXK,TSPAN5,MT1E,TM4SF1,IFFO2,TNFRSF10D,C11orf68,SPHK1,KIRREL1,S100A3,PLXNB3,MICB,KRTAP2-3,LYN |
| ETS1-FOSL1 | repressed | SEMA3F,USH1C,IL20RA,CNTN1,SYT13,GYG2,REEP1,MAP2,CECR2,NOL4,SCG3,FZD3,NCALD,TLE6,CHN2,TMEM176B,DDX25,ADGRD1,HMGCS1,IGFBP2,SGIP1,HSPA2,TSPAN8,PIK3C2B,NTS,CMPK2,FXYD6,STXBP5L,CHRNB2,ELAVL4,TMEM169,NDRG2,CRABP1,RAB26,NPNT,ENHO,PARM1,GTSF1,GPR27,KCNK3,PCSK1,CRYBG2,RIMKLA,MKRN3,NRXN1,IDH2,PCP4,RIPPLY3,SBK1,TMEM198,KCNMB2,CASTOR3,SELENOP,CBSL |
| ETS1-SMAD3 | activated | MRC2,ANLN,LIMA1,DCBLD2,IFI35,ACSL4,CD59,GADD45B,SMURF2,CPZ,KLHL5,GNAI2,CLIP4,UAP1,GBP1,IFIT3,TNFRSF10B,RASSF8,ACOT9,SDC4,TRIP10,ARMCX1,CALU,STARD13,IL6ST,RAB11FIP5,FAM129A,DDX60,IFI44,FRMD6,COL6A1,EFHD2,SH3BGRL3,CYR61,ABL2,NCEH1,LHFPL2,ARHGAP18,SH3KBP1,UGCG,TAGLN,CDC42EP2,RHOC,ZYX,ITGA5,NEXN,CDC42EP3,IFI16,PTX3,ANXA5,EDIL3,GPX8,CGAS,ABCA1,LETM2,SERPINB8,SLFN5,TPM4,PXDC1,NINJ2,RIN1,OXTR,F2R,PRR16,KCNQ5,TRABD2A,PDCD1LG2,SAMD9,APOBEC3G,PRSS2 |
| ETS1-SMAD3 | repressed | TMEM176A,ABCC8,CACNA2D2,NLRP2,CDH1,SYNE2,KCNH2,SEZ6,STAG3,LLGL2,CEACAM6,NRCAM,TRIM9,TOX3,NCALD,CAPS,PTPRZ1,GLCCI1,CPE,SIPA1L2,PRXL2A,POF1B,DLL4,APOE,GADD45G,GCH1,DDC,SYT4,FAM83F,UNC79,BICDL1,CGA,SULF1,TTC6,AK7,CCDC33,RNF165,RGS16,N4BP3,CPLX2,ADGRL3,AKR1C2,RNF144A,MPV17L,BTG2,CELF3,SPTBN4,WNT4,VSNL1,TMEM169,WNK2,AQP3,PHYHIPL,SEC11C,DEGS2,MFSD2A,AGR3,MFSD4A,CALCB,PRR15,TCIM,SRRM3,GLDC,PCP4,MAP7D2,PLA2G6,KCND2,RPS27L,C12orf56,CYP4F3,CARMIL3,RASSF10,SFMBT2,RBM20,UBE2QL1,CKMT1B,MYO15B |
| FOSL1-SMAD3 | activated | CYP26B1,COPZ2,FHL1,TMSB10,ADAMTS6,CYBA,SYNJ2,CD40,TIMP1,KLHL4,SLC15A3,HBEGF,GNAI2,IFI27L2,PPP1R3C,SERPINB6,ATXN1,CPA4,C19orf66,FST,OASL,CALHM2,CDH11,COL6A2,TM4SF19,LYPD1,GJA1,ANGPT1,SH3RF2,IL31RA,LARP6,NNMT,CAVIN3,PRNP,SPTLC3,EFEMP2,TUBB6,MXRA7,COL18A1,EVI2B,PLEKHG4,FADS3,GPM6B |
| FOSL1-SMAD3 | repressed | EYA2,SPTB,ST6GALNAC2,SULT2B1,SLC4A11,CHGA,NCALD,SPOCK2,TBC1D30,EYA4,RTKN,PIK3R3,AIF1L,FGD3,TNFRSF19,MAP7,SEMA6D,SLC44A3,CGN,INA,TMEM45B,CACNA1D,TNFRSF14,RADIL,TMEM169,TMEM184A,CLDN3,COL22A1,NRG4,PCP4,ABAT,UTY,KLRG2,DENND1C,FAM19A5,PNMA6A |
| ETS1-FOSL1-SMAD3 | activated | GNAI2 |
| ETS1-FOSL1-SMAD3 | repressed | NCALD,TMEM169,PCP4 |

**Supplementary Table. S4** Gene Ontology Enrichment for the differentially expressed target genes

| ONTOLOGY | Description | GeneRatio | BgRatio | pvalue | p.adjust | qvalue | Count |
|---|---|---|---|---|---|---|---|
| **HALLMARKS** | EPITHELIAL MESENCHYMAL TRANSITION | 0,1224 | 183/3376 | 7,650E-22 | 3,825E-20 | 2,738E-20 | 107 |
| | KRAS SIGNALING UP | 0,0881 | 161/3376 | 6,892E-10 | 1,723E-08 | 1,233E-08 | 77 |
| | APICAL JUNCTION | 0,0824 | 159/3376 | 4,732E-08 | 6,582E-07 | 4,711E-07 | 72 |
| | TNFA SIGNALING VIA NFKB | 0,0870 | 171/3376 | 5,265E-08 | 6,582E-07 | 4,711E-07 | 76 |
| | INFLAMMATORY RESPONSE | 0,0744 | 146/3376 | 4,786E-07 | 4,786E-06 | 3,426E-06 | 65 |
| | HYPOXIA | 0,0767 | 171/3376 | 6,308E-05 | 5,257E-04 | 3,763E-04 | 67 |
| | IL2 STAT5 SIGNALING | 0,0664 | 155/3376 | 8,006E-04 | 4,448E-03 | 3,184E-03 | 58 |
| | ESTROGEN RESPONSE EARLY | 0,0732 | 177/3376 | 1,240E-03 | 5,637E-03 | 4,035E-03 | 64 |
| | ESTROGEN RESPONSE LATE | 0,0686 | 167/3376 | 2,088E-03 | 8,701E-03 | 6,228E-03 | 60 |
| **BIOLOGICAL PROCESS** | CELL JUNCTION ORGANIZATION | 0,0847 | 498/11194 | 9,528E-23 | 4,062E-19 | 3,226E-19 | 197 |
| | CELL PART MORPHOGENESIS | 0,0744 | 497/11194 | 8,339E-14 | 3,555E-11 | 2,824E-11 | 173 |
| | SYNAPTIC SIGNALING | 0,0731 | 461/11194 | 3,260E-16 | 4,632E-13 | 3,680E-13 | 170 |
| | REGULATION OF ION TRANSPORT | 0,0684 | 430/11194 | 2,380E-15 | 2,537E-12 | 2,015E-12 | 159 |
| | TRANSMEMBRANE RECEPTOR PROTEIN TYROSINE KINASE SIGNALING PATHWAY | 0,0675 | 434/11194 | 3,210E-14 | 1,890E-11 | 1,501E-11 | 157 |
| | CELL MORPHOGENESIS INVOLVED IN NEURON DIFFERENTIATION | 0,0658 | 420/11194 | 3,546E-14 | 1,890E-11 | 1,501E-11 | 153 |
| | POSITIVE REGULATION OF LOCOMOTION | 0,0645 | 402/11194 | 6,333E-15 | 5,400E-12 | 4,289E-12 | 150 |
| | TAXIS | 0,0641 | 418/11194 | 5,805E-13 | 2,062E-10 | 1,638E-10 | 149 |
| | BLOOD VESSEL MORPHOGENESIS | 0,0632 | 428/11194 | 2,126E-11 | 5,035E-09 | 4,000E-09 | 147 |
| | RESPONSE TO GROWTH FACTOR | 0,0632 | 468/11194 | 2,284E-08 | 2,562E-06 | 2,035E-06 | 147 |
| **MOLECULAR FUNCTIONS** | CALCIUM ION BINDING | 0,0779 | 453/10453 | 2,624E-16 | 2,134E-13 | 1,696E-13 | 165 |
| | ION TRANSMEMBRANE TRANSPORTER ACTIVITY | 0,0694 | 497/10453 | 2,655E-07 | 1,349E-05 | 1,072E-05 | 147 |
| | CELL ADHESION MOLECULE BINDING | 0,0652 | 433/10453 | 4,112E-09 | 3,714E-07 | 2,953E-07 | 138 |
| | STRUCTURAL MOLECULE ACTIVITY | 0,0595 | 493/10453 | 2,054E-03 | 2,037E-02 | 1,619E-02 | 126 |
| | INORGANIC MOLECULAR ENTITY TRANSMEMBRANE TRANSPORTER ACTIVITY | 0,0567 | 413/10453 | 8,750E-06 | 2,156E-04 | 1,714E-04 | 120 |
| | ACTIN BINDING | 0,0562 | 339/10453 | 7,293E-11 | 1,976E-08 | 1,571E-08 | 119 |
| | PHOSPHOLIPID BINDING | 0,0562 | 345/10453 | 2,577E-10 | 4,322E-08 | 3,436E-08 | 119 |
| | SIGNALING RECEPTOR REGULATOR ACTIVITY | 0,0538 | 329/10453 | 4,535E-10 | 5,267E-08 | 4,187E-08 | 114 |
| **REACTOME** | NERVOUS SYSTEM DEVELOPMENT | 0,0878 | 452/7427 | 1,226E-06 | 8,710E-05 | 7,565E-05 | 135 |
| | HEMOSTASIS | 0,0872 | 440/7427 | 4,057E-07 | 3,364E-05 | 2,922E-05 | 134 |
| | SIGNALING BY RECEPTOR TYROSINE KINASES | 0,0820 | 386/7427 | 1,140E-08 | 1,890E-06 | 1,642E-06 | 126 |
| | RHO GTPASE CYCLE | 0,0768 | 361/7427 | 3,139E-08 | 3,905E-06 | 3,391E-06 | 118 |
| | SIGNALING BY GPCR | 0,0755 | 404/7427 | 5,072E-05 | 1,577E-03 | 1,370E-03 | 116 |
| | EXTRACELLULAR MATRIX ORGANIZATION | 0,0683 | 236/7427 | 4,974E-17 | 4,950E-14 | 4,299E-14 | 105 |

**Supplementary Table. S5** Mean Influence (MI) of regulators in DMSO, Capmatinib or Trametinib treatment under HGF stimulation conditions

| Regulators | DMSO+HGF | | CAPM+HGF | | TRAM+HGF | |
|---|---|---|---|---|---|---|
| | MI | SD | MI | SD | MI | SD |
| PPARG | 2,42 | 0,67 | 1,37 | 0,37 | -2,62 | 0,45 |
| STAT6 | -0,7 | 0,19 | 0,66 | 0,99 | -0,51 | 0,2 |
| YAP1 | -0,88 | 0,5 | 0,68 | 0,63 | -0,17 | 0,9 |
| TCF4 | -1,87 | 0,21 | -0,64 | 0,74 | 2,22 | 0,36 |
| TWIST2 | 0,4 | 0,32 | 0,6 | 1,09 | -0,48 | 0,31 |
| MAGEA1 | -0,49 | 0,27 | 0,51 | 0,4 | -1,13 | 0,59 |
| MSC | 2,84 | 0,69 | -1,93 | 1,76 | -0,07 | 0,99 |
| AUTS2 | -0,88 | 0,53 | 0,13 | 0,34 | 0,63 | 0,43 |
| OVOL2 | -1,91 | 0,26 | -2,42 | 0,67 | 2,5 | 0,21 |
| DACH1 | -2,25 | 0,49 | -0,09 | 0,59 | 1,55 | 0,81 |
| SOX2 | 0,88 | 0,08 | 0,08 | 0,83 | -1,48 | 0,45 |
| TESC | -0,71 | 0,66 | -2,11 | 1,7 | 1,73 | 2,46 |
| EGFR | -0,82 | 0,22 | 0,16 | 0,61 | 0,4 | 0,41 |
| FHL2 | -1,12 | 0,39 | 0,08 | 0,29 | 0,32 | 0,3 |
| LMO2 | 0,83 | 0,33 | -0,19 | 0,47 | -0,3 | 0,81 |
| NKX2-5 | 0,6 | 0,54 | 0,77 | 0,37 | -0,96 | 0,8 |
| PTPN14 | 0,48 | 0,25 | 0,42 | 0,84 | -0,74 | 0,14 |
| HES6 | -1,44 | 0,62 | -1,42 | 0,37 | 2,46 | 0,49 |
| SIX2 | 1,55 | 0,22 | 0,57 | 0,46 | -2,1 | 0,31 |
| ST18 | -1,74 | 0,31 | -0,87 | 0,52 | 0,92 | 0,51 |
| CBX2 | -0,69 | 0,46 | -0,05 | 0,71 | 1,16 | 0,54 |
| FOSL1 | 3,18 | 0,54 | -0,46 | 0,73 | -1,78 | 0,32 |
| SMAD3 | 4,81 | 0,48 | -0,66 | 0,44 | -2,98 | 0,44 |
| INSM1 | -0,46 | 0,59 | -1,41 | 1,16 | 0,97 | 0,31 |
| MYB | -0,06 | 0,78 | 1,29 | 0,37 | -0,42 | 0,61 |
| MNX1 | 2,39 | 0,3 | -0,11 | 0,77 | -0,81 | 0,11 |
| NOTCH2 | 0,01 | 0,87 | -2,06 | 1,01 | 1,82 | 0,43 |
| ETS1 | 1,93 | 0,9 | -2,64 | 0,86 | 0,67 | 0,67 |
| FOXO6 | 0,61 | 0,23 | -0,41 | 0,96 | -0,06 | 0,39 |
| PSIP1 | -1,81 | 0,48 | -1,15 | 0,7 | 1,96 | 0,24 |
| ZNF711 | -2,03 | 0,27 | -0,31 | 1,08 | 1,98 | 0,6 |
| CBFA2T2 | -0,85 | 0,2 | -0,25 | 0,47 | 0,92 | 0,42 |
| KMT5C | -0,81 | 0,71 | -1,7 | 0,81 | 1,88 | 0,51 |
| SETBP1 | -1,2 | 0,35 | -1,15 | 1,4 | 1,78 | 0,31 |
| IRF6 | -0,96 | 0,35 | -0,06 | 0,56 | 0,26 | 0,41 |
| OVOL1 | 0,62 | 0,51 | 0,02 | 0,63 | -0,84 | 0,42 |
| PAX5 | -1,17 | 0,58 | 0,64 | 1,07 | 0,32 | 0,46 |
| ZNF3 | -1,29 | 0,45 | -1,61 | 0,59 | 2,45 | 0,69 |
| TGFB1I1 | 0,1 | 0,28 | -1,18 | 0,81 | 0,41 | 0,22 |
| ZFP30 | -0,5 | 0,38 | 1,17 | 0,48 | -0,63 | 0,76 |
| ZNF620 | -0,59 | 0,27 | 0,03 | 0,45 | 0,33 | 0,11 |
| ZNF853 | -0,4 | 0,14 | -0,58 | 0,47 | 0,89 | 0,64 |
| LOXL2 | -3,31 | 0,89 | -0,17 | 0,49 | 1,57 | 0,22 |
| DLX5 | 0,03 | 0,17 | 1,14 | 0,55 | -1,44 | 0,35 |
| ACTL6B | -0,22 | 0,35 | 1,24 | 0,75 | 0,09 | 0,27 |
| MYT1 | 1,47 | 0,86 | 0,05 | 0,79 | -1,12 | 0,3 |
| HOXD10 | -1,32 | 0,62 | -0,22 | 0,62 | 0,62 | 0,5 |
| NR0B1 | -0,87 | 0,34 | 0,06 | 0,83 | 0,73 | 0,08 |
| SYK | -1,69 | 0,65 | -0,19 | 1,12 | 0,99 | 0,14 |
| CCND1 | 4,02 | 0,71 | 2 | 0,61 | -3,87 | 0,55 |
| NFKB2 | -0,53 | 0,79 | 1,31 | 0,47 | -1,37 | 0,41 |
| FOXA1 | -1,43 | 0,15 | 0,79 | 0,49 | 1,14 | 0,32 |
| NKX2-1 | -1,39 | 0,24 | 0,69 | 0,27 | 1,01 | 0,09 |
| LMNA | -0,27 | 0,34 | -0,4 | 0,97 | 0,32 | 0,42 |
| GRHL2 | -0,46 | 0,27 | -0,63 | 1 | -0,08 | 0,28 |
| NOTCH3 | -0,4 | 0,55 | -0,99 | 0,64 | 0,55 | 0,24 |
| FGFR2 | -0,4 | 0,74 | 0,72 | 0,92 | -1,39 | 1,01 |
| TWIST1 | -1,65 | 0,48 | -1,2 | 0,79 | 1,85 | 0,52 |
| PROX1 | 0,4 | 0,29 | -0,79 | 0,27 | 0,39 | 0,32 |
| CAMK4 | 0,69 | 0,37 | 0,84 | 1,16 | 0,1 | 0,89 |
| DLX2 | -0,1 | 0,14 | 0,19 | 0,86 | -0,14 | 0,26 |
| FOXG1 | 0,26 | 0,25 | 0,2 | 0,42 | -0,46 | 0,5 |
| HOXD11 | 1,07 | 0,46 | -0,07 | 0,94 | -0,81 | 0,22 |
| ZNF397 | -1,08 | 0,25 | -0,34 | 0,48 | 0,62 | 0,36 |
| NR4A1 | 2,42 | 1,02 | 0,67 | 0,72 | -2,3 | 2,9 |
| SALL2 | -1,13 | 0,5 | 0,33 | 0,36 | 0,95 | 0,33 |
| ZNF775 | -0,35 | 0,21 | 0,6 | 0,38 | -0,05 | 0,53 |
| MEIS3 | -0,45 | 0,36 | 0,02 | 1,59 | 0,37 | 0,7 |
| TBX3 | -0,06 | 0,69 | 2,85 | 1,55 | -1,12 | 1,02 |
| BEX1 | 0,55 | 0,36 | 0,25 | 0,91 | -0,64 | 0,49 |
| CDK5R1 | 0,89 | 0,33 | 0,34 | 0,41 | -0,48 | 0,17 |
| E2F2 | -0,8 | 0,42 | -1,22 | 0,36 | 1,23 | 0,12 |
| HOXA4 | 1,14 | 0,23 | 0,07 | 0,69 | -0,18 | 0,29 |
| POU4F1 | 1 | 0,24 | 0,65 | 1 | -0,77 | 0,14 |
| ZNF708 | 0,31 | 0,67 | 0,89 | 0,9 | -0,89 | 0,19 |
| ZNF135 | 0,08 | 0,72 | -0,01 | 1,22 | 0,17 | 0,13 |
| ZNF804A | -0,1 | 0,19 | -0,12 | 0,22 | 0,16 | 0,14 |
| MYO6 | -0,76 | 0,54 | -0,77 | 0,67 | 0,74 | 0,6 |
| ZFP3 | -1,5 | 0,37 | -2,01 | 1,01 | 1,29 | 0,28 |

| Regulators | DMSO+HGF | | CAPM+HGF | | TRAM+HGF | |
|---|---|---|---|---|---|---|
| | MI | SD | MI | SD | MI | SD |
| WWTR1 | 2,89 | 0,29 | 0,23 | 0,68 | -2,29 | 0,42 |
| FOS | 1,25 | 0,56 | 0,14 | 0,62 | -1,03 | 0,41 |
| ONECUT2 | -1,18 | 0,19 | -0,34 | 0,65 | 1,09 | 0,23 |
| ISL1 | -0,76 | 0,64 | 0,21 | 0,56 | 0,1 | 0,14 |
| RUNX1 | 0,01 | 0,88 | 0,57 | 0,9 | 0,15 | 0,3 |
| DLX6 | -1,53 | 0,22 | 0,26 | 0,7 | 1,24 | 0,39 |
| HOXD13 | 0,62 | 0,26 | -0,13 | 1,25 | -0,28 | 1,18 |
| TEAD3 | -0,51 | 0,55 | -1,67 | 2,96 | 1,04 | 0,43 |
| TGFB1 | 2,99 | 0,27 | 0,91 | 0,26 | -2,51 | 0,18 |
| AJUBA | -1,56 | 0,81 | -0,46 | 0,66 | 0,56 | 0,37 |
| MAML3 | 0,89 | 0,44 | -0,76 | 0,48 | 0,07 | 0,37 |
| FOSB | 0,99 | 0,3 | 0,08 | 0,89 | -1,05 | 0,55 |
| ZNF512 | -1,34 | 0,48 | -0,17 | 0,6 | 0,67 | 0,41 |
| HES1 | 0,35 | 0,49 | -1,19 | 0,33 | 0,28 | 0,71 |
| MAGEA2 | -1,18 | 0,24 | 0,05 | 1,04 | 1,05 | 0,54 |
| CLU | -2,32 | 1,22 | -0,85 | 1,58 | 1,4 | 0,43 |
| DAB2 | 0,44 | 0,48 | -0,5 | 0,68 | 0,29 | 0,33 |
| KLF5 | -1,03 | 0,29 | 0,38 | 0,41 | -0,09 | 0,34 |
| SOX4 | -0,44 | 0,28 | 0,09 | 0,94 | 0 | 0,22 |
| CHD7 | 0,04 | 0,23 | 0,39 | 0,4 | -0,2 | 0,23 |
| ELK3 | -0,18 | 0,19 | 0,38 | 0,35 | -0,07 | 0,45 |
| HMGA2 | 7,33 | 0,78 | 2,72 | 0,63 | -5,88 | 0,61 |
| EHF | -0,9 | 0,3 | 0,48 | 1,05 | 0,42 | 0,19 |
| ELF3 | -0,87 | 0,44 | -0,4 | 1,26 | 0,29 | 0,4 |
| AHR | -0,79 | 0,38 | 0,08 | 0,29 | 0,01 | 0,19 |
| TLE4 | -2,28 | 3,8 | 0,49 | 0,89 | -0,06 | 1,18 |
| ZNF821 | -2,43 | 0,71 | 0,41 | 1,06 | 1,07 | 0,36 |
| RELB | -1,89 | 0,5 | 1,37 | 1,28 | -0,2 | 0,44 |
| CAVIN1 | -2,2 | 0,72 | -0,38 | 0,23 | 1,26 | 0,62 |
| MECOM | 2,83 | 1,13 | -0,22 | 0,32 | -2,45 | 0,51 |
| APBB1 | 0,81 | 0,68 | 0,11 | 0,66 | -0,95 | 1,2 |
| ENG | 1,93 | 0,58 | -1,04 | 1,15 | -1,19 | 0,53 |
| ZNF93 | -0,95 | 0,48 | 0,63 | 0,64 | -0,1 | 1,63 |
| ZEB1 | 1,5 | 0,44 | -1,29 | 1,37 | -0,1 | 0,24 |
| MITF | 0,69 | 0,39 | -0,65 | 0,07 | -0,23 | 0,24 |
| GLI3 | -0,72 | 0,05 | -0,9 | 1,06 | 0,94 | 0,29 |
| POU3F2 | -1,25 | 0,58 | -0,48 | 0,4 | 1,18 | 0,31 |
| ZEB2 | 0,63 | 0,35 | 0,21 | 0,75 | -0,78 | 0,13 |
| ZIC1 | 0,14 | 0,26 | 1,49 | 0,79 | -0,74 | 0,48 |
| PAX6 | 1,77 | 0,41 | -0,17 | 1,2 | -0,95 | 0,79 |
| CASZ1 | -1,47 | 0,65 | -0,09 | 0,3 | 1,28 | 0,21 |
| H2AFY2 | 0,88 | 0,56 | 0,96 | 0,22 | -0,83 | 0,27 |
| ASCL1 | 0,11 | 0,27 | 0,72 | 0,87 | -0,35 | 0,55 |
| EGR1 | 0,71 | 0,59 | 1,09 | 0,66 | -0,91 | 0,55 |
| JUP | 1,33 | 0,21 | 0 | 0,72 | -1,47 | 0,35 |
| FLNA | -0,91 | 1,1 | 2,03 | 3,18 | -1,14 | 1,11 |
| TP73 | 0,75 | 0,88 | 0,56 | 1 | -0,42 | 0,52 |
| BHLHE40 | 2,29 | 0,45 | 0,94 | 0,31 | -2,73 | 0,31 |
| TBX1 | 1,77 | 0,27 | -3,31 | 2,28 | 0,29 | 0,44 |
| RUNX2 | 1,4 | 0,11 | 0,44 | 0,44 | -0,98 | 0,14 |
| SMAD9 | 0,19 | 0,37 | -0,07 | 0,38 | -0,18 | 0,7 |
| BARX2 | -0,84 | 0,63 | 0,36 | 1,49 | 2,59 | 2,43 |
| MUC1 | -1,37 | 0,15 | -0,63 | 0,55 | 1,2 | 0,24 |
| ARHGEF5 | -1,83 | 0,55 | 0,65 | 1,2 | 0,81 | 0,34 |
| RARG | -2,1 | 0,22 | -0,65 | 0,61 | 1,32 | 0,89 |
| HOXC10 | 1,01 | 0,48 | -0,6 | 1,13 | -0,6 | 0,42 |
| HOXC13 | -0,19 | 0,55 | -1,03 | 1,16 | 1,21 | 0,58 |
| HOXC6 | -0,59 | 0,67 | 0,1 | 0,39 | 0,54 | 0,58 |
| ZNF493 | -0,88 | 0,73 | 1,85 | 1,64 | -1,01 | 0,39 |
| SP100 | -0,21 | 0,68 | 0,52 | 1,05 | -0,57 | 0,48 |
| FGF2 | -2,25 | 0,49 | -0,44 | 0,89 | 1,94 | 0,32 |
| HOXD9 | 0,77 | 0,36 | 0,49 | 0,39 | -0,27 | 0,47 |
| BCOR | 0,01 | 0,29 | -0,7 | 0,22 | 0,46 | 0,12 |
| EPAS1 | -2,64 | 0,52 | 0,64 | 0,48 | 0,51 | 0,48 |
| TFAP2A | -1,13 | 0,48 | 0,09 | 0,41 | 0,2 | 0,06 |
| BNC1 | 0,2 | 0,17 | 1,27 | 0,83 | -0,04 | 0,29 |
| HOXC9 | 0,35 | 0,36 | 0,62 | 0,76 | -0,28 | 0,66 |
| TSHZ1 | -1,04 | 0,27 | -1,47 | 0,34 | 1,4 | 0,14 |
| IFI27 | -1,89 | 0,08 | -0,71 | 0,62 | 0,95 | 0,26 |
| NPAS2 | 0,35 | 0,15 | -0,11 | 0,59 | -0,56 | 0,32 |
| VDR | -0,69 | 0,39 | -0,69 | 0,28 | 1,17 | 0,41 |
| SNAI2 | -1,98 | 0,36 | 0,88 | 0,46 | 0,33 | 0,27 |
| ATF7IP2 | -1,58 | 0,13 | 0,89 | 1,55 | 0,51 | 0,77 |
| ZNF257 | 2,34 | 0,48 | 1,03 | 0,55 | -2,16 | 0,32 |
| GATA4 | 0,15 | 0,37 | 0,71 | 0,38 | -0,98 | 0,37 |
| SP4 | 0,18 | 0,31 | 0,62 | 0,53 | -1,25 | 0,43 |
| TDG | 0,98 | 0,36 | 1,1 | 0,52 | -1,03 | 0,3 |
| AEBP1 | -1,74 | 1 | 0 | 0,45 | 0,81 | 1,35 |
| CXXC4 | 0,67 | 0,25 | 1,72 | 1,36 | -1,87 | 0,58 |

| Regulators | DMSO+HGF | | CAPM+HGF | | TRAM+HGF | |
|---|---|---|---|---|---|---|
| | MI | SD | MI | SD | MI | SD |
| HSF2 | -1,69 | 0,39 | -1,07 | 0,95 | 1,99 | 0,53 |
| LHX2 | 0,14 | 0,25 | 0,21 | 0,71 | -0,36 | 0,38 |
| SNCA | -0,17 | 0,3 | -1,95 | 1,42 | 1,28 | 0,5 |
| ZNF704 | 0,59 | 0,2 | -0,3 | 0,33 | -0,06 | 0,36 |
| ZIK1 | -1,27 | 0,1 | -0,62 | 0,18 | 0,8 | 0,17 |
| FOSL2 | 1,89 | 1,15 | -1,97 | 1,46 | -0,06 | 0,4 |
| MYCL | 0,92 | 0,25 | 0,56 | 0,25 | -1,39 | 0,25 |
| PRMT6 | 0,14 | 0,54 | 1,8 | 1,09 | -1,21 | 0,49 |
| PITX2 | -0,97 | 0,35 | -1,62 | 1,57 | 2,89 | 1,32 |
| CEBPB | 1,18 | 0,22 | 0,34 | 0,92 | -1,39 | 1,27 |
| ZNF195 | 1,66 | 0,89 | 2,45 | 2,46 | -2,77 | 0,46 |
| ESRRG | 0,12 | 0,51 | 0,66 | 0,88 | -0,5 | 0,5 |
| IRX2 | 1,46 | 0,97 | 0,58 | 0,15 | -0,63 | 0,26 |
| IRF9 | -1,71 | 1,24 | 1,18 | 1,15 | 0,25 | 0,78 |
| PHF8 | 0,39 | 0,24 | -1,13 | 0,57 | 1,36 | 0,25 |
| CREB3L4 | -0,8 | 0,3 | -1,61 | 0,77 | 1,04 | 0,19 |
| DZIP1 | -0,57 | 0,44 | 1,15 | 0,47 | -0,6 | 0,96 |
| NFIX | 0,29 | 0,13 | 0,47 | 0,31 | -0,14 | 0,31 |
| ZNF608 | -0,17 | 0,17 | -0,5 | 0,21 | 0,71 | 0,15 |
| DUSP26 | 0,96 | 0,21 | -0,19 | 0,44 | -0,53 | 0,55 |
| PLCB1 | 1,79 | 0,27 | 1,13 | 0,67 | -2,51 | 0,46 |
| RBPMS | 0,25 | 0,47 | -0,38 | 0,91 | -0,17 | 0,17 |
| ARNT2 | -0,76 | 0,41 | -0,01 | 0,29 | 0,69 | 0,56 |
| JUNB | 0,02 | 0,47 | 0,25 | 1,93 | -0,3 | 0,32 |
| SOX21 | 0,07 | 0,62 | 0,25 | 0,1 | 0,1 | 0,26 |
| ZFP69B | -0,07 | 0,3 | 0,38 | 0,8 | -0,07 | 0,18 |
| BARX1 | -0,14 | 0,2 | 0,07 | 0,6 | -0,32 | 0,45 |
| DPF1 | 0,42 | 0,41 | 0,18 | 0,48 | -0,2 | 0,55 |
| CRTC1 | -2,03 | 0,43 | -1,5 | 0,69 | 1,67 | 0,37 |
| LEF1 | 2,1 | 0,76 | -0,12 | 0,45 | -1,14 | 0,88 |
| CBFA2T3 | -2,06 | 0,71 | 1,16 | 1,2 | 0,7 | 0,75 |
| MSX2 | 0,06 | 0,36 | -1,24 | 1,31 | 0,71 | 1,3 |
| CITED1 | 1,03 | 0,23 | 0,06 | 0,8 | -0,93 | 0,47 |
| VGLL1 | 1,99 | 0,44 | 0,74 | 0,89 | -1,74 | 0,89 |
| MYT1L | 1,06 | 0,42 | -0,51 | 0,13 | -0,32 | 0,44 |
| HOXB7 | -0,99 | 1,18 | -0,59 | 0,17 | 0,99 | 0,71 |
| ZBED2 | 0,08 | 0,95 | -0,94 | 0,35 | 1,4 | 0,71 |
| TFEB | -1,41 | 0,3 | 0,01 | 1,32 | 1,58 | 0,75 |
| HOXC8 | 0,66 | 0,72 | 0,09 | 1,97 | 0,74 | 1,15 |
| H3F3A | -0,92 | 0,25 | 0,81 | 0,29 | 0,01 | 0,27 |
| BCL9L | 0,12 | 0,64 | 1,07 | 0,78 | -0,42 | 0,58 |
| DLX1 | -0,72 | 0,65 | -1,29 | 1,36 | 0,84 | 0,58 |
| FOXP2 | -1,4 | 1 | -1,81 | 1,51 | 2,07 | 1,31 |
| GFI1 | 0,74 | 0,37 | 0,63 | 0,16 | -0,85 | 0,38 |
| TSPYL2 | -0,78 | 0,25 | 1 | 0,72 | -0,07 | 0,14 |
| PRAME | 0,77 | 0,72 | 2,16 | 0,95 | -2,62 | 1,45 |
| HOXB8 | 0,22 | 0,62 | -0,6 | 1,78 | 1,12 | 0,87 |
| HOXD8 | 0,15 | 0,34 | 2,5 | 1,47 | -0,75 | 0,5 |
| ZNF43 | -2,39 | 1,33 | -0,07 | 0,64 | 1,28 | 0,31 |
| S100A9 | 0,57 | 0,37 | -0,29 | 1,66 | 0,43 | 0,6 |
| TNKS | 0,95 | 0,22 | -0,16 | 0,75 | -1,02 | 0,69 |
| FOXA2 | 0,48 | 0,39 | 0,2 | 0,6 | -0,37 | 0,37 |
| TLE2 | -0,46 | 0,38 | 1,65 | 0,67 | -0,29 | 0,37 |
| HOXA9 | 1,27 | 0,56 | 0,54 | 0,99 | -0,56 | 1,01 |
| REPIN1 | 0,54 | 0,25 | -0,69 | 0,51 | -0,06 | 0,22 |
| MAGEA2B | -1,57 | 0,33 | -2,27 | 0,32 | 2,15 | 0,43 |
| FLI1 | -0,24 | 0,6 | -0,3 | 0,65 | 0,43 | 0,86 |
| TFAP2C | -0,99 | 0,84 | 1,32 | 0,83 | -0,66 | 0,25 |
| SALL1 | 1,34 | 0,19 | 0,2 | 0,43 | -0,95 | 0,19 |
| ETV4 | 3,01 | 0,1 | 1,84 | 0,66 | -2,3 | 0,18 |
| POU2F2 | 1,7 | 0,46 | 2,62 | 0,95 | -2,82 | 0,84 |
| ZNF664 | 0,22 | 0,27 | 1,43 | 1,2 | -0,78 | 0,77 |
| ZSCAN30 | 0,07 | 0,5 | 1,33 | 0,67 | -0,96 | 0,34 |
| GLIS3 | -0,36 | 0,19 | -0,27 | 0,22 | 0,42 | 0,12 |
| HMGA1 | 0,79 | 0,62 | 0,82 | 1,21 | -0,94 | 0,78 |
| FOXQ1 | 1,27 | 0,67 | 1,59 | 1,54 | -0,75 | 0,82 |
| TBX18 | 0,3 | 0,42 | -0,32 | 0,77 | 0,01 | 0,35 |
| TRIM29 | 0,93 | 0,26 | 0,19 | 0,85 | -1,61 | 0,88 |
| GATA3 | 0,57 | 0,6 | 0,73 | 0,34 | -0,52 | 0,35 |
| JDP2 | -2,1 | 1,31 | -3,42 | 4,79 | 1,42 | 1,31 |
| DLX3 | -1,74 | 0,55 | 1,23 | 1,17 | 0,59 | 0,52 |
| SIX3 | 0,85 | 0,64 | -0,68 | 1,02 | 0,48 | 0,51 |
| FRK | 0,8 | 0,67 | -1,57 | 0,79 | 0,72 | 0,55 |
| SOX11 | 0,89 | 0,4 | 1,52 | 0,89 | -1,64 | 0,4 |
| NR4A2 | 0,72 | 1,28 | 0,51 | 1,08 | -1,86 | 0,37 |

| Regulators | DMSO+HGF | | CAPM+HGF | | TRAM+HGF | |
|---|---|---|---|---|---|---|
| | MI | SD | MI | SD | MI | SD |
| DRAP1 | 1,52 | 0,42 | -0,44 | 0,76 | -0,97 | 0,44 |
| GRHL1 | 0,11 | 0,7 | -0,65 | 1,32 | -0,07 | 0,52 |
| HOXA10 | -0,98 | 0,5 | 0,78 | 0,66 | -0,31 | 0,29 |
| IRAK2 | -0,83 | 0,34 | 0,23 | 0,31 | 0,71 | 0,33 |
| SP110 | -0,66 | 0,4 | 0,6 | 1 | 0,08 | 0,88 |
| HOXB3 | -0,68 | 0,65 | -0,59 | 1,05 | 0,71 | 0,44 |
| EGLN3 | -0,22 | 0,44 | -0,11 | 0,32 | 0,31 | 0,62 |
| EYA1 | 1,26 | 0,2 | 1,02 | 0,29 | -1,13 | 0,16 |
| TGIF2 | 1,16 | 0,34 | 0,21 | 0,6 | -0,61 | 0,37 |
| HNF1B | 0,82 | 0,09 | 1,18 | 0,17 | -1,18 | 0,07 |
| LOXL3 | -0,52 | 0,14 | -0,81 | 0,55 | 0,58 | 0,28 |
| POU6F2 | -1,6 | 1,4 | 0,95 | 0,81 | -1,14 | 0,93 |
| ZNF677 | -1,24 | 0,53 | -0,36 | 0,3 | 2,18 | 0,53 |
| TRIM5 | -2,32 | 0,75 | 0,07 | 0,3 | 5,88 | 6,8 |
| PBX1 | 0,99 | 0,29 | 0,99 | 1,79 | -1,65 | 0,56 |
| SKAP1 | -1,09 | 0,35 | 0,59 | 0,91 | 0,54 | 1,26 |
| SPDEF | 1,07 | 0,38 | -1,33 | 0,47 | 0,18 | 0,42 |
| TFCP2L1 | 0,78 | 0,35 | 0,94 | 0,63 | -0,76 | 1,04 |
| ZNF300 | -2,2 | 4,89 | -0,29 | 0,28 | 0,2 | 0,23 |
| BASP1 | -0,34 | 0,28 | 0,68 | 0,26 | -0,12 | 0,31 |
| CRABP2 | -1,26 | 0,43 | -1,5 | 0,21 | 1,97 | 0,29 |
| ZFY | -1,32 | 0,34 | -0,09 | 0,49 | 0,74 | 0,12 |
| ZNF667 | -0,28 | 1,26 | -0,23 | 0,62 | 1,17 | 1,66 |
| IKZF2 | -1,38 | 0,4 | -1,9 | 0,5 | 1,81 | 0,68 |
| ZNF528 | -0,94 | 0,16 | -1,31 | 0,5 | 1,23 | 0,18 |
| BATF | -0,9 | 0,15 | -1,17 | 0,86 | 1,73 | 0,23 |
| RORC | 0,35 | 0,06 | 0,83 | 0,48 | -0,47 | 0,12 |
| ZNF516 | -0,31 | 0,38 | -1,43 | 0,91 | 0,71 | 0,39 |
| S100A8 | 0,73 | 0,27 | 0,59 | 0,9 | -0,53 | 0,39 |
| L3MBTL4 | -0,81 | 1,92 | 1,33 | 1,15 | -1,17 | 0,7 |
| EBF4 | -0,82 | 0,29 | 0,85 | 0,64 | -0,22 | 0,77 |
| KLF6 | -1,36 | 0,18 | 2,09 | 1,16 | 0,37 | 0,37 |
| NRG1 | 2,18 | 0,56 | 0,26 | 0,35 | -1,4 | 0,57 |
| IRF1 | -8,27 | 14,73 | -0,1 | 0,67 | 0,97 | 0,47 |
| SGK1 | -1,68 | 0,26 | -1,84 | 0,53 | 1,36 | 0,18 |
| HOXD4 | 1,19 | 0,28 | -0,44 | 0,7 | -0,86 | 0,41 |
| PID1 | 0,81 | 0,23 | 0,65 | 0,55 | -0,58 | 0,22 |
| ETV1 | -1,66 | 0,64 | -1,34 | 0,53 | 1,44 | 1,43 |
| CDKN2B | -1,04 | 0,41 | -0,73 | 0,37 | 0,95 | 0,53 |
| ZNF714 | -1,06 | 0,23 | -0,28 | 2,14 | 0,72 | 0,32 |
| ZNF334 | -1,34 | 0,97 | -0,21 | 0,41 | 1,03 | 0,57 |
| SOX7 | 0,34 | 0,23 | -0,85 | 0,25 | 0,71 | 0,18 |
| NFATC4 | -1,13 | 0,28 | -0,54 | 0,66 | 1,28 | 0,38 |
| TRIM22 | -0,16 | 0,46 | 0,11 | 0,7 | 0,36 | 0,33 |
| ZNF559 | -0,76 | 0,32 | -1,79 | 0,81 | 1,47 | 0,36 |
| HOXB2 | 0,16 | 0,94 | 0,32 | 0,78 | -0,83 | 0,55 |
| FOXF2 | -2,52 | 0,64 | 1,4 | 0,41 | -1 | 0,26 |
| PARP10 | -1,08 | 0,57 | 0,64 | 0,72 | -0,26 | 0,57 |
| RUNX3 | -0,53 | 0,4 | 0,51 | 0,98 | 0,24 | 0,4 |
| HEYL | -0,43 | 1,1 | -0,41 | 0,79 | 0,79 | 0,91 |
| PHF21A | -1,99 | 0,89 | 1,19 | 0,66 | -0,23 | 0,64 |
| MAD2L2 | 0,15 | 0,55 | 0,06 | 0,9 | -0,07 | 0,37 |
| CCDC85B | 1,71 | 0,38 | -0,52 | 0,38 | -0,85 | 0,16 |
| ZSCAN18 | -1,51 | 0,79 | 0,12 | 0,62 | 1,35 | 0,52 |
| ATF5 | 1,94 | 0,83 | 1,88 | 0,73 | -2,25 | 0,47 |
| SOX13 | -2,07 | 0,82 | -1,07 | 1,23 | 1,37 | 0,78 |
| NMI | 1,65 | 0,34 | 0,12 | 0,56 | -1,3 | 0,31 |
| ZNF439 | -2,14 | 0,5 | 1,67 | 0,87 | 0,4 | 0,34 |
| SIM2 | 0,28 | 0,31 | 1,96 | 1,06 | -0,56 | 0,3 |
| HOXA7 | -0,33 | 0,45 | -1,14 | 1,04 | 1,69 | 1,58 |
| ZNF532 | -0,73 | 0,88 | 0,53 | 0,46 | -0,06 | 0,8 |
| HIPK2 | -0,39 | 0,19 | -1,54 | 0,79 | 0,86 | 0,16 |
| PIR | -0,01 | 0,41 | -0,81 | 0,32 | 0,4 | 0,14 |
| HOXD3 | 0,27 | 0,18 | -0,05 | 0,74 | 0,21 | 0,27 |
| MESP1 | 0,64 | 0,27 | -0,94 | 0,28 | 0,57 | 0,23 |
| PITX1 | 0,13 | 0,79 | 0,77 | 0,91 | -0,62 | 0,36 |
| ZNF513 | 0,27 | 0,17 | 0,92 | 1,1 | -0,17 | 0,33 |
| YBX2 | -0,52 | 0,29 | 0,81 | 0,2 | -0,56 | 0,33 |
| ZNF682 | 0,1 | 0,12 | -0,52 | 0,45 | 0,4 | 0,29 |
| ZNF626 | -0,38 | 1,03 | -0,4 | 1,15 | 0,21 | 0,74 |
| NFE2L3 | 2,15 | 0,63 | -0,04 | 1,13 | -1,71 | 1,65 |
| HOXB3 | -0,68 | 0,65 | -0,59 | 1,05 | 0,71 | 0,44 |
| EGLN3 | -0,22 | 0,44 | -0,11 | 0,32 | 0,31 | 0,62 |
| EYA1 | 1,26 | 0,2 | 1,02 | 0,29 | -1,13 | 0,16 |
| TGIF2 | 1,16 | 0,34 | 0,21 | 0,6 | -0,61 | 0,37 |
